# PCRD-seq: Proximity Crosslinking-induced RNA Depletion for Low-Input Subcellular Transcriptome Profiling

**DOI:** 10.64898/2026.01.17.700046

**Authors:** Jinghua Han, Lan Li, Xueqing Kong, Kaiyun Zhao, Yuen Ting Leung, Alice S. T. Wong, Ying Li

## Abstract

Chromatin-associated RNAs play critical roles in regulating chromatin organization and transcription, underscoring the importance of their study. Proximity labeling has emerged as a promising and versatile technique for profiling chromatin-associated RNAs with high spatiotemporal resolution. While being a powerful technique, traditional proximity labeling methods depend on complex, high-input enrichment protocols, which significantly limit their wide practical application. Here, we developed a straightforward, enrichment-free chromatin-associated RNA profiling strategy: Proximity Crosslinking-induced RNA Depletion sequencing (PCRD-seq). This approach leverages the proximity crosslinking between chromatin and its surrounding RNAs induced by singlet oxygen generated by HoeDBF, a photosensitizer targeting chromatin region. The proximity crosslinking hinders the release of chromatin-associated RNAs during routine TRIzol extraction, consequently leading to a specific depletion of these RNAs. This method was successfully applied to investigate the role of U1 snRNA in RNA chromatin retention and the differences in chromatin-associated transcriptomes between two ovarian cancer cell lines with opposite metastatic capability. Moreover, our PCRD-seq exhibits potential in profiling nuclear lamina-associated RNAs, which paves the way for its application to profile RNAs associated with other chromatin subdomains. The minimal cell input and simple workflow endow PCRD-seq as a transformative tool for wide applications.

## Introduction

In eukaryotes, RNAs are asymmetrically distributed and their precise subcellular localization is fundamental to their function ^1–3^. Among these localized pools, chromatin-associated RNAs (caRNAs) have emerged as key regulatory molecules, playing vital and direct roles in shaping chromatin architecture, modulating transcriptional programs, and contributing to epigenetic memory, thereby influencing essential cellular states and pathways ^4, 5^.

Mapping caRNAs is therefore critical, and several strategies have been developed. Classical approaches rely on subcellular fractionation to isolate chromatin pellet, from which caRNAs are extracted for sequencing ^6, 7^. To profile RNAs associated with specific chromatin states, immunoprecipitation-based approaches, such as Chromatin RNA immunoprecipitation followed by sequencing (ChRIP-seq) ^8^, Profiling interacting RNAs on chromatin followed by deep sequencing (PIRCh-seq) ^9^, and Chromatin-associated RNA immunoprecipitation followed by next-generation sequencing (CARIP-Seq) ^10^, use antibodies to purify crosslinked chromatin bearing desired epigenetic marks, followed by isolation and sequencing of the co-purified RNAs. Reverse Transcribe and Tagment (RT&Tag) detects RNAs at specific genomic loci via antibody-targeted *in situ* reverse transcription and tagmentation of the resulting RNA-cDNA hybrid. Alternatively, proximity ligation methods like Mapping RNA-genome interactions (MARGI) ^11^, Chromatin-associated RNA sequencing (ChAR-seq) ^12^ and Global RNA interactions with DNA by deep sequencing (GRID-seq) ^13^ capture genome-wide RNA-DNA interactions by ligating crosslinked RNA-DNA pairs with specialized linkers.

Despite being powerful, these techniques have limitations. Fractionation can induce contamination, while the other methods, which are antibody- or crosslinking-dependent, may not faithfully capture native, live-cell interactions and suffer from high background as well as antibody batch variability ^14^. Furthermore, protocols for proximity ligation are particularly complex and time-consuming, imposing a significant practical burden ^12, 13^.

Proximity labeling (PL) has emerged as a promising alternative, enabling the tagging of biomolecules in living cells under near-native conditions ^15^. This technique utilizes engineered enzymes, photocatalysts or photosensitizers to generate reactive species for the labeling of proximal molecules, a strategy that has been successfully applied to profile caRNAs in nuclear bodies ^16^ and at specific epigenetic marks ^14^. We previously contributed to this toolkit with two photosensitizer-based systems: 1) the Halo-DBF system in which a photosensitizer dibromofluorescein (DBF) is directed to specific subcellular locations via HaloTag system for proximity labeling ^17^; 2) the small molecule HoeDBF ^18^, which utilizes a Hoechst moiety to target DBF to chromatin directly for caRNA profiling ^19^. In these systems, light-activated DBF produces singlet oxygen (^1^O_2_), which oxidizes the guanosines on proximal RNAs to highly reactive intermediates OG^ox^. The OG^ox^ reacts with propargylamine (PA), by which an alkyne handle is installed onto the OG^ox^. The alkynated sites are further biotinylated via Cu(I)-catalyzed azide-alkyne cycloaddition (CuAAC) reaction ^20^ for streptavidin-based enrichment.

However, almost all PL methods, including our own, rely on multi-step biotin enrichment. This process is cumbersome, imposes a significant burden on sample preparation and introduces technical variability due to complex handling. Most critically, the enrichment-based methods require high RNA input, posing a substantial barrier for low-input applications. The development of enrichment-free PL methodologies is poised to advance the field drastically. Recently, ONIC-seq ^21^ emerged as the first and only such methodology. It profiles localized RNAs via detecting mutations induced by guanine oxidation. However, this method involves a complex pipeline for the detection of guanine mutations.

Inspired by meCLICK-seq, a method for profiling m^6^A-modified RNAs through specific depletion of these transcripts ^22^, we hypothesized that caRNAs could similarly be detected through targeted depletion, utilizing a standard differential expression analysis pipeline. According to established literature showing that oxidized guanine in DNA can form crosslinks with proximal proteins ^23, 24, 25^, we hypothesized that OG^ox^ intermediates on RNA could analogously react with chromatin proteins during DBF-mediated PL at chromatin. This process would create RNA-protein-DNA crosslinks, selectively trapping caRNAs on chromatin and impeding their release during standard TRIzol extraction. The resultant depletion of caRNAs in the soluble RNA fraction thus provides a direct, enrichment-free readout.

Here, we developed an enrichment-free caRNA profiling strategy, termed Proximity Crosslinking-induced RNA Depletion Sequencing (PCRD-seq). Based on small molecule HoeDBF system as well as genetically encoded HaloTag system, PCRD-seq leverages ^1^O_2_-induced crosslinking to profile caRNAs through their specific depletion, overcoming the key limitations of enrichment-based protocols. We applied PCRD-seq to investigate U1 snRNA-mediate RNA chromatin retention and to identify caRNA signatures distinguishing ovarian cancer cell lines with opposite metastatic potential, demonstrating its versatile utility in addressing biological questions.

## Results

### HoeDBF-mediated proximity labeling specially captures caRNAs

First, we validated the sensitivity and specificity of HoeDBF-mediated proximity labeling using fluorescence confocal imaging, cellular fractionation coupled with RNA dot blot analysis, and enrichment followed by qRT-PCR (Fig. 1A). The chromatin localization of HoeDBF is mediated by its Hoechst moiety. This was supported by live-cell imaging, where co-staining with Hoechst 33342 competitively displaced HoeDBF, resulting in a decreased and diffuse DBF signal (Fig. S1A). Therefore, Hoechst 33342 counterstaining was omitted for HoeDBF-treated cells in live-cell imaging. Despite this, HoeDBF fluorescence revealed well-defined nuclear shapes (Fig. 1B), indicating its effective nuclear localization. Fixed-cell fluorescence imaging of labeled biomolecules further validated their nuclear enrichment (Fig. 1C and Fig. S1B). To determine if the labeling was specific to caRNAs, we performed cell fractionation followed by RNA dot blot assay. Biotin signals, representing labeled RNAs, were primarily detected in the chromatin fraction, with minimal signal in the nucleoplasm and negligible in the cytoplasm (Fig. 1D). Finally, quantitative real time polymerase chain reaction (qRT-PCR) analysis of biotinylated RNAs enriched by streptavidin-pulldown demonstrated specific enrichment of known chromatin-associated transcripts (*XIST*, *MALAT1*, *DDX5*) in HoeDBF-labeled cells, but not cytosolic markers (*GAPDH* and *ACTB*) (Fig. 1E). Collectively, these results demonstrated that HoeDBF-mediated proximity labeling specially captures caRNAs.

**Fig. 1.**
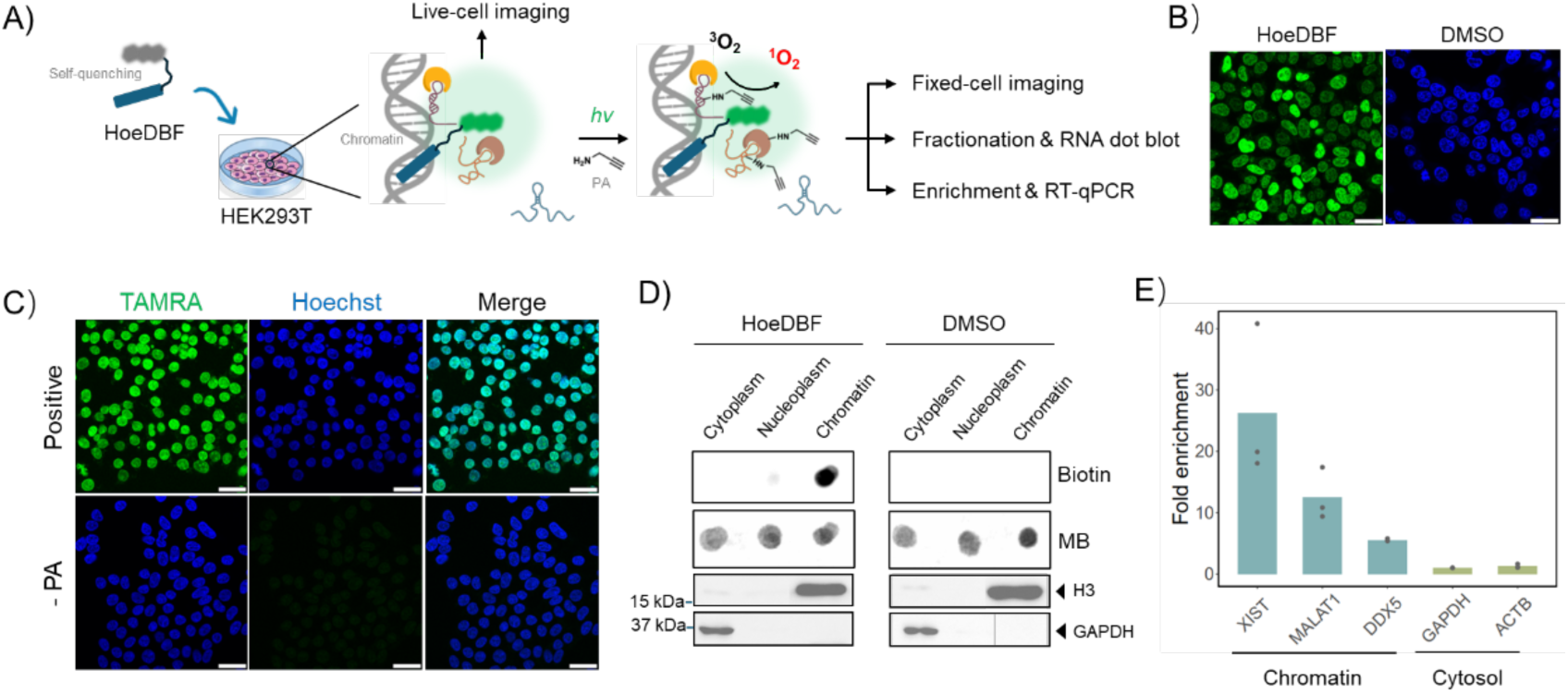
The specificity and sensitivity of HoeDBF-mediated proximity labeling. (A) Schematic diagram illustrating the experimental design for the validation of specificity and sensitivity of HoeDBF-mediated proximity labeling. (B) Fluorescence confocal imaging in living cells after incubation with HoeDBF. Green signal represents HoeDBF molecules bound to chromatin. Blue signal represents Hoechst 33342 nucleic acid stain. Nuclei were not counterstained for HoeDBF treated cells to avoid the displacement of HoeDBF by Hoechst 33342. Scale bar: 25 μm. (C) Fluorescence confocal imaging in fixed cells after HoeDBF labeling. TAMRA was ligated to labeled biomolecules for visualization. Scale bar: 25 μm. (D) The figures in the upper two rows indicate RNA dot blot results of different cellular locations after HoeDBF labeling and cell fractionation. The figures in the lower two rows indicate western blot results of the chromatin marker protein H3 and cytosolic marker protein GAPDH. The signal of biotin represents labeled RNAs. MB: methylene blue. DMSO treated cells were used as negative control. (E) qRT-PCR results showing the fold enrichment of chromatin and cytosol marker genes.

### HoeDBF-mediated proximity labeling enables transcriptome-wide profiling of caRNAs

Having established the sensitivity and specificity of HoeDBF labeling, we performed transcriptome-wide profiling of caRNAs in HEK293T cells by integrating the enrichment protocol with RNA-seq (Fig. 2A). RNAs significantly up-regulated in enriched versus input samples (log_2_ (fold change) ≥ 1 and FDR < 0.05) were defined as caRNAs, yielding 3,372 candidates, including well-documented examples such as *XIST*, *KCNQ1OT1*, *PVT1*, *MALAT1*, and *TSIX* (Fig. 2B).

**Fig. 2.**
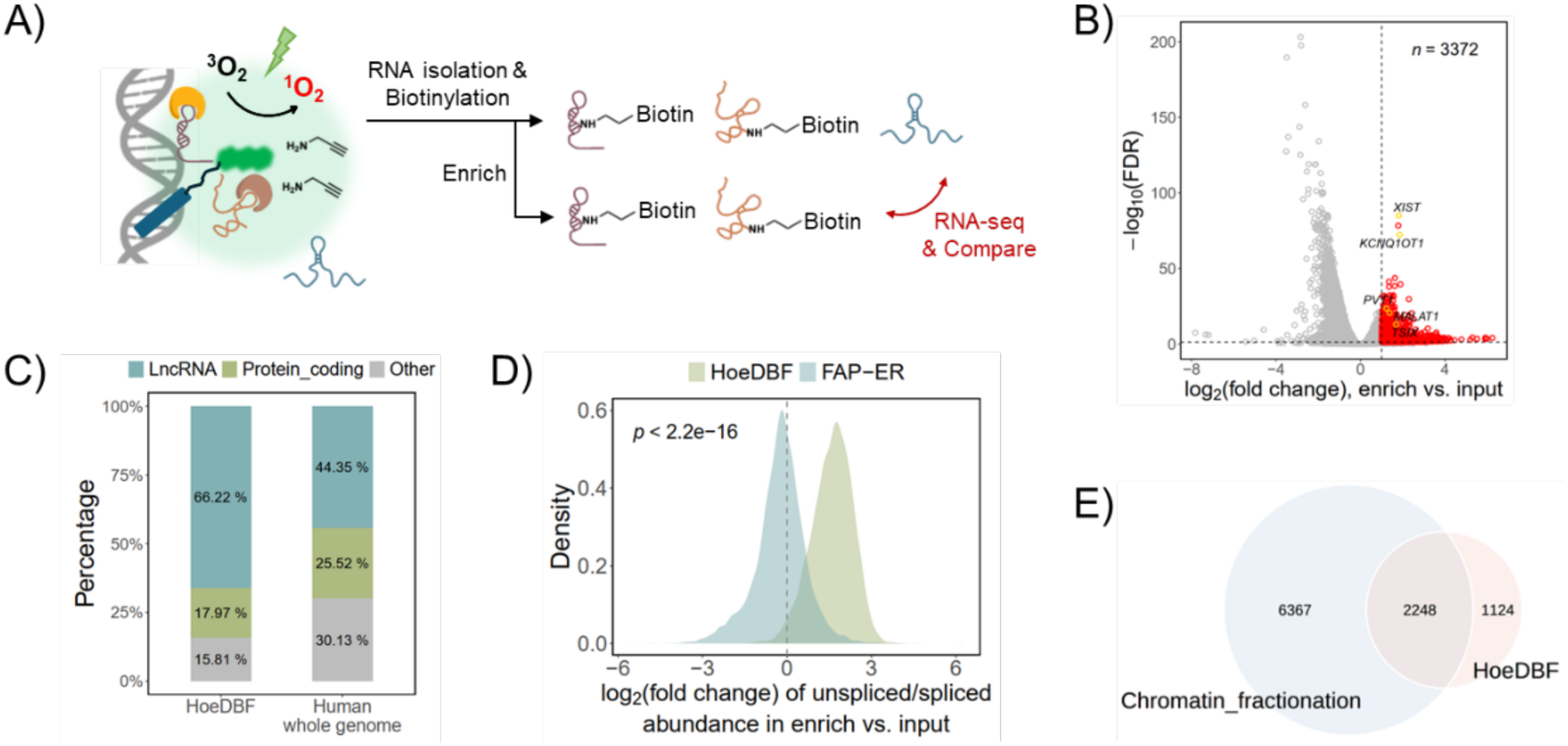
Transcriptomic profiling of the caRNAs in HEK293T cells using HoeDBF-mediated proximity labeling. (A) Schematic illustration of the key experimental procedures. (B) Volcano plot showing the differential expression between enriched and input samples. Red dots indicate significantly up-regulated genes in enriched samples, i.e., caRNAs. Five well-documented caRNAs were highlighted in yellow. (C) Gene biotype compositions of the caRNAs identified by HoeDBF-mediated proximity labeling and human whole genome. (D) The distribution of the log_2_(fold change) of the unspliced-to-spliced ratio in pulldown versus input samples. The unspliced-to-spliced ratio indicates the ratio of unspliced to spliced isoform abundance for each transcript. The RNA-seq data of our FAP-ER proximity labeling system was used for comparison. The grey dashed line corresponds to log_2_(fold change) = 0, indicating that the unspliced-to-spliced ratios in pulldown samples are equal to those in the input. Differences were analyzed using Wilcoxon test. (E) Venn diagram showing the overlap between the caRNAs identified by HoeDBF-mediated proximity labeling and the reported chromatin fractionation approach.

We next explored the characteristics of these caRNAs in detail. Biotype analysis revealed a strong enrichment for lncRNAs (66.22%) compared to the human whole genome (44.35%) (Fig. 2C), consistent with the reported nature of caRNAs ^5, 26^. As RNAs are transcribed from chromatin and spliced in nuclear speckles ^27, 28^, we hypothesized that enriched samples would contain a higher proportion of unspliced transcripts compared to input samples. Indeed, the unspliced-to-spliced transcript ratios were markedly higher in enriched samples (Fig. 2D, green curve), aligning well with previous report ^29^. As a control, we analyzed the data from our FAP-ER proximity labeling system for the profiling of endoplasmic reticulum (ER)-localized RNAs, which are primarily mature mRNAs ^30^. As expected, the unspliced-to-spliced ratios were not elevated in FAP-ER enriched samples (Fig. 2D, blue curve). This contrast confirms that HoeDBF method captures RNAs at chromatin. Finally, comparison with a published chromatin fractionation dataset ^7^ showed that two-thirds of our identified caRNAs were also detected by this orthogonal method (Fig. 2E). Together, these results established that HoeDBF-mediated proximity labeling enables transcriptome-wide identification of caRNAs with high confidence.

### Development of an enrichment-free caRNA profiling strategy (PCRD-seq)

Conventional enrichment-based proximity labeling (PL) strategies, including our initial HoeDBF method, are subject to technical variabilities. A clear example is the sensitivity of caRNA identification to the washing stringency during streptavidin-biotin enrichment, which substantially altered our results (Fig. S2A). To circumvent these protocol-dependent variabilities, we devised an enrichment-free alternative: Proximity Crosslinking-induced RNA Depletion sequencing (PCRD-seq). We hypothesized that ^1^O_2_ generated during HoeDBF labeling could induce covalent crosslinks between chromatin and nearby RNAs, thereby trapping caRNAs and impeding their release during TRIzol extraction. This specific depletion could then be quantified by sequencing, forming the basis of PCRD-seq (Fig. 3A).

**Fig. 3.**
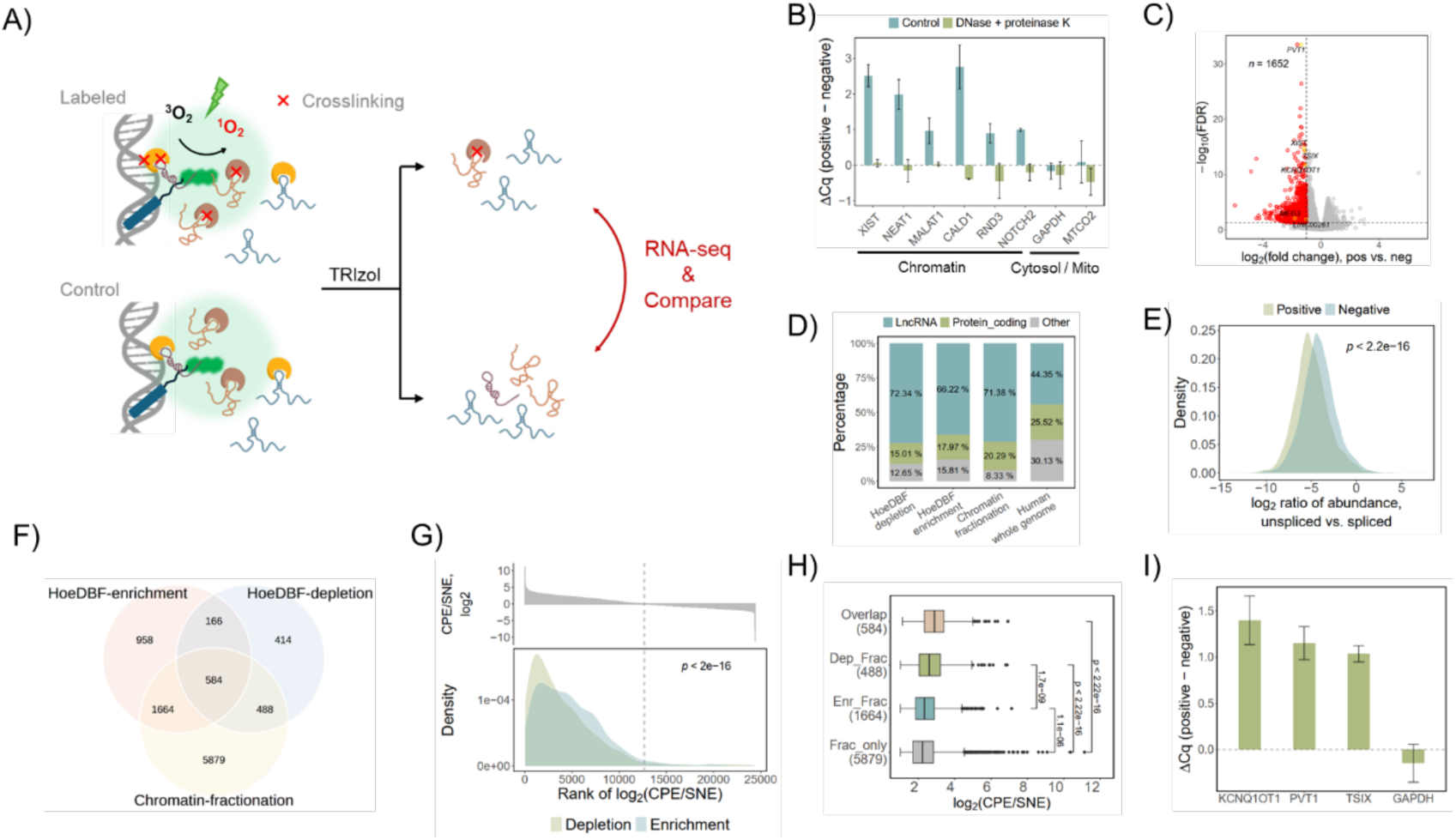
PCRD-seq enables the transcriptome-wide profiling of caRNAs. (A) Illustration of the principle of PCRD-seq. (B) qRT-PCR analysis of RNAs isolated from DNase and Proteinase K treated cell lysate. (C) Volcano plot showing the differential expression between HoeDBF labeled (positive) and control (negative) cells. Red dots indicate transcripts that are significantly down-regulated in positive group (log_2_(fold change) ≤ -1, FDR < 0.05), i.e., caRNAs. Six well-documented caRNAs were highlighted in yellow. (D) Gene biotype composition of caRNAs identified by different approaches. The result of human whole genome serves as background. (E) Density plot showing the distribution of log_2_-transformed unspliced transcript enrichment factor in positive and negative groups. The unspliced transcript enrichment factor was calculated as the ratio of unspliced-to-spliced isoform abundance for each transcript. Differences were analyzed using Wilcoxon test. (F) Venn diagram showing the overlaps of caRNAs identified by different approaches. (G) The x-axis of this figure indicates the rank of caRNAs identified in chromatin fractionation approach based on their log_2_(CPE/SNE) values. The bar plot in the upper panel shows the log_2_(CPE/SNE) values of these ranked caRNAs. The density plot in the lower panel shows the distribution of the ranking positions for caRNAs identified in our depletion- or enrichment-based approach. The grey dashed line indicates the ranking position of the caRNA with the smallest positive log_2_(CPE/SNE) value (1.37e-05). Differences were analyzed using Wilcoxon test. (H) The log_2_(CPE/SNE) values for different caRNA sets showed in (F). (I) The ΔCq values (positive vs. negative) for the caRNAs identified by PCRD-seq and cytosolic maker gene GAPDH.

We first verified that ^1^O_2_ exposure during HoeDBF labeling specifically induces the depletion of caRNAs. qRT-PCR showed that HoeDBF labeling significantly increased the Cq values (indicating depletion) for chromatin marker genes, but not for cytosolic or mitochondrial markers, compared to the negative control (Fig. 3B and Fig. S2B). To confirm that this observation was caused by RNA-protein-DNA crosslinking, we treated cell lysates with DNase and Proteinase K to digest crosslinked adducts. This enzymatic treatment abolished the difference in Cq values for chromatin markers between HoeDBF-labeled cells and the negative control (Fig. 3B), validating that ^1^O_2_-induced RNA-chromatin crosslinking is the underlying mechanism for the observed RNA depletion.

We next optimized labeling conditions by testing two regimes expected to generate weak (5 μM HoeDBF, 2-min light) or strong (10 μM HoeDBF, 3-min light) crosslinking, each with two negative controls (no light or no HoeDBF), affording total four pairs of conditions for comparison: “5 μM, no *hv*”, “5 μM, no HoeDBF”, “10 μM, no *hv*” and “10 μM, no HoeDBF” (Tabel S1). CCK-8 assays revealed 85.3% and 64.3% cell viability for the weak and strong labeling conditions, respectively (Fig. S2C). Furthermore, we noticed much higher cytotoxicity for 10 μM HoeDBF incubation even without green light irradiation (87.7%) compared to 5 μM (99%; Fig. S2C). Using criteria of log_2_(fold change) ≤ -1 and FDR < 0.05, RNA-seq analysis identified comparable numbers of depleted RNAs (caRNAs) under the weak and strong labeling conditions, irrespective of the paired negative controls (Figs. S2D, S2E and S2F). However, their overlap was poor (Figs. S2G and S2H), a discrepancy we ascribed to the pronounced cytotoxicity of 10 μM HoeDBF. For negative control selection, since RNAs were isolated immediately after light exposure, before potential light-induced influence could manifest, we reasoned that the “no light” control best normalizes for the influence of HoeDBF incubation. Based on these findings, we selected the “5 μM, no *hv*” condition for all subsequent studies.

Applying this optimized PCRD-seq protocol, we identified 1,652 potential caRNAs, including some well-documented chromatin marker genes (Figs. 3C and S3). The biotype composition analysis demonstrated that the caRNAs detected using the depletion-based approach were predominantly lncRNAs, with an even higher proportion than those detected in our enrichment-based approach and the reported chromatin fractionation-based method ^7^ (Fig. 3D). Moreover, consistent with the expectation that unspliced chromatin-associated transcripts would be crosslinked and depleted, we observed a lower unspliced-to-spliced ratio in labeled samples versus negative controls (Fig. 3E). Finally, a substantial fraction (45.4% and 64.9%) of these caRNAs overlapped with those identified by our enrichment-based method and the fractionation-based approach ^7^, respectively (Fig. 3F). Taking all these results together, we conclude that PCRD-seq enables the capture of high-confidence caRNAs.

We hypothesized that PCRD-seq preferentially detects RNAs with stronger chromatin association compared to enrichment-based method. This hypothesis may partly explain the observation that the number of caRNAs identified by PCRD-seq is much lower than that identified by enrichment-base method. To test this, we used the log_2_(CPE/SNE) values from the fractionation study as a proxy for chromatin-association strength ^7^. CPE and SNE represent chromatin pellet extract and soluble nuclear extract, respectively, and the CPE/SNE ratio indicates the relative abundance of RNA between these two fractions. caRNAs detected via depletion-based approach ranked significantly higher than those detected by enrichment-based approach according to their log_2_(CPE/SNE) values (Fig. 3G). Consistently, caRNAs uniquely identified by depletion- and fractionation-base methods (“Dep_Frac”) showed significantly higher log_2_(CPE/SNE) values than those specific to enrichment- and fractionation-based methods (“Enr_Frac”; Fig. 3H). These findings confirm that PCRD-seq enriches for caRNAs with tighter chromatin associations. Finally, qRT-PCR independently validated the depletion of three identified caRNAs (*KCNQ1OT1*, *PVT1*, and *TSIX*; Fig. 3I).

In summary, we developed PCRD-seq, a straightforward method that profiles caRNAs via proximity crosslinking-induced depletion and preferentially captures RNAs with high chromatin-association strength.

### Application of PCRD-seq in other subcellular locations

To test the versatility of PCRD-seq, we applied it to profile RNAs associated with nuclear lamina, a filamentous protein network at the inner nuclear membrane ^31^ that interacts with peripheral heterochromatin via lamina-associated domains (LADs) to modulate chromatin architecture ^32, 33^. To target DBF to nuclear lamina, we utilized HEK293 cells stably expressing a Halo-LMNA fusion protein. A total of 2,009 candidates were identified (Figs. 4A and 4B), most of which were lncRNAs (67.15%; Fig. 4C). Approximately two-thirds were also detected by either the HoeDBF-mediated enrichment- or depletion-based approaches (Fig. 4D), suggesting their association with chromatin. Despite the substantial overlap with HoeDBF-captured caRNAs, which may be due to the nucleoplasmic distribution of some Lamin A/C ^34, 35^, PCRD-seq in Halo-LMNA-expressing cells demonstrated higher specificity in detecting lamina-associated RNAs. This was evidenced by the enrichment of nuclear pore complex interacting protein family members among the top highest-confidence protein-coding RNA hits (6 out of the top 11; Fig. 4B), a signature absent from HoeDBF-captured caRNAs (Fig. S3). Taken together, these results demonstrated the capability of PCRD-seq in characterizing lamina-associated RNAs and paved the way for its application to profile RNAs associated with other chromatin subdomains, such as histone modification sites or transcription factor binding sites.

**Fig. 4.**
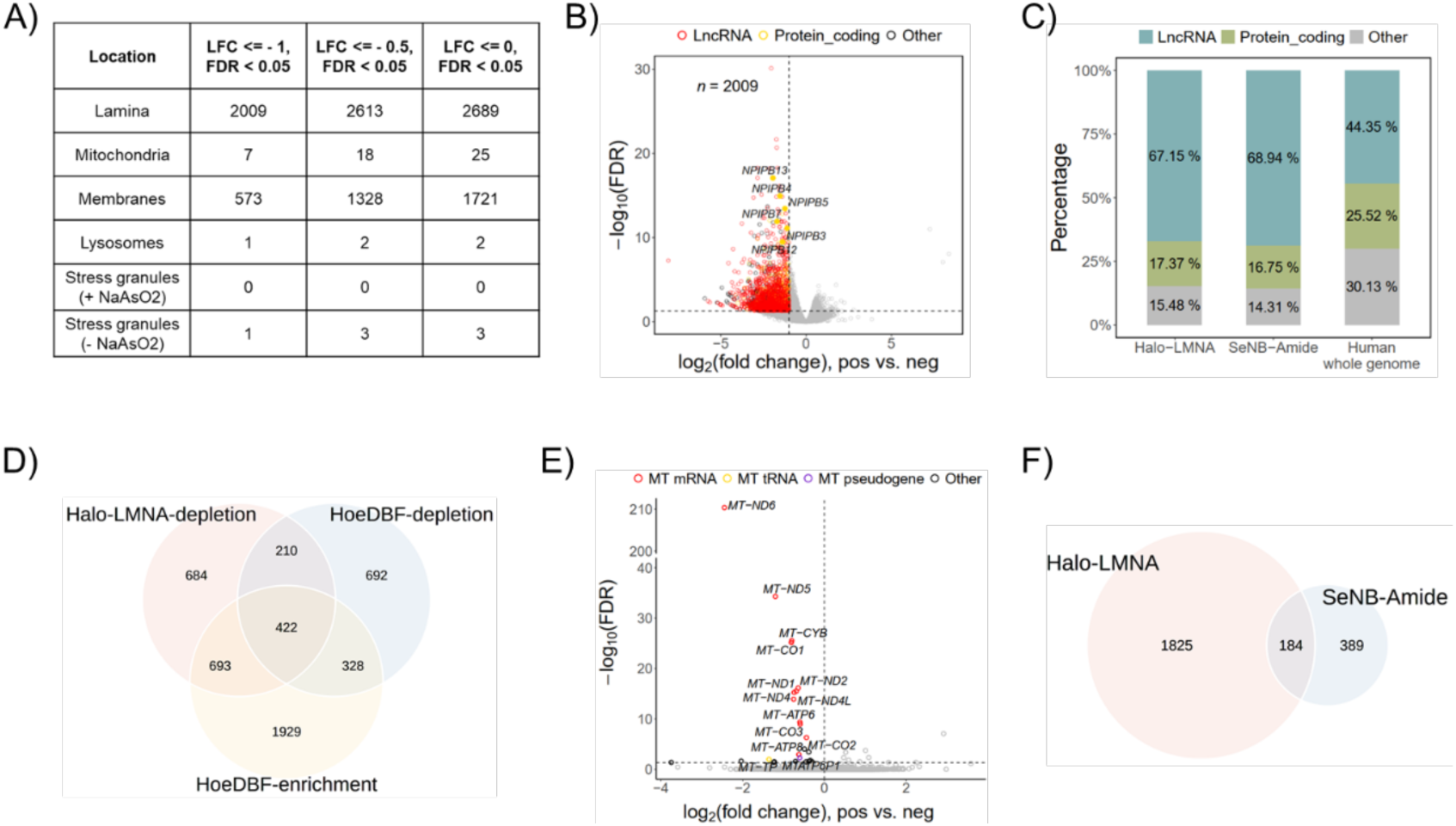
(A) The number of RNAs localized in different cellular compartments identified using different log_2_(fold change) (LFC) cutoffs. (B) Volcano plot showing the differential expression between labeled (positive) and control (negative) Halo-LMNA-expressing cells. Red and yellow dots indicate significantly down-regulated lncRNAs and protein coding RNAs (LFC ≤ -1, FDR < 0.05), respectively, i.e., lamina-associated RNAs. The yellow solid dots highlight the nuclear pore complex interacting protein family members among the top highest-confidence protein-coding RNA hits. (C) Gene biotype compositions of lamina-associated RNAs, endomembrane-associated RNAs and human whole genome. (D) Venn diagram showing the overlaps between lamina-associated RNAs and caRNAs identified by HoeDBF-mediated enrichment-and depletion-based approaches. (E) volcano plot showing the differential expression between SeNB-Mito labeled (positive) and control (negative) U2OS cells. Red, yellow and purple dots represent mitochondrial mRNAs (MT mRNA), tRNAs (MT tRNA) and pseudogene (MT pseudogene), respectively. Black dots represent other differentially expressed transcripts. The horizontal grey dashed line denotes - log_10_(FDR) = 1.301, while the vertical grey dashed line represents the threshold of LFC = 0. (F) Venn diagram showing the overlap between lamina-associated RNAs (Halo-LMNA) and endomembrane-associated RNAs (SeNB-Amide).

We next investigated whether proximity crosslinking-induced RNA depletion is generalizable to other subcellular locations. Using small-molecule probes that target endomembranes (mainly ER, SeNB-Amide), mitochondria (SeNB-Mito) and lysosomes (SeNB-Lyso) ^36^, we performed PCRD-seq across these locations. Since these small-molecule probes were originally developed and validated in U2OS cells, all related experiments in this study were consequently performed in the same cell line to ensure optimal performance. To profile stress granule-associated RNAs, we employed our previously established FAP-MGHI PL system ^37^. Specifically, we utilized a HeLa cell line stably expressing FAP-G3BP1 fusion protein and applied sodium arsenite to induce stress granule formation. Significant RNA depletion was specifically detected for mitochondria and endomembranes (Fig. 4A). The mitochondria dataset showed high-confidence depletion of 13 mitochondrial genome-encoded RNAs (12 mRNAs, 1 tRNA) and 1 mitochondrial pseudogene encoded by nuclear genome, while other hits had markedly lower confidence (Fig. 4E). The identified endomembrane-associated RNAs were predominantly lncRNAs (68.94%; Fig. 4C) and approximately one-third overlapped with our identified lamina-associated RNAs (Fig. 4F), likely reflecting SeNB-Amide’s labeling of nuclear envelope. We conclude that proximity crosslinking-induced RNA depletion is particularly effective in DNA-rich environments (chromatin, mitochondrial genome) or at DNA-proximal membranes, where DNA may act as a scaffold for crosslinking complexes.

Collectively, these results establish PCRD-seq as a simple yet versatile tool for profiling RNAs associated with chromatin, its subdomains, and the mitochondrial genome.

### Investigating the role of U1 snRNA in RNA chromatin retention using PCRD-seq

U1 snRNP has been implicated in lncRNA chromatin retention in mouse embryonic stem cells (mESCs) ^38^. We used our HoeDBF-mediated PCRD-seq to test if this mechanism is conserved in human cells. We inhibited U1 snRNA in HEK293T cells using antisense morpholino oligonucleotides (AMOs) to block its 5’-end recognition sequence. Next, we employed our PCRD-seq to characterize the alterations in the chromatin-associated transcriptome following U1 snRNA inhibition (Fig. 5A). To evaluate the efficiency of U1 snRNA blocking, we established a method that integrates RNase H protection assay with qRT-PCR ^39^ (Fig. S4A). This method was first validated using total RNA (Fig. S4B). Next, we conducted U1 AMO concentration titration experiment to determine the optimal concentration. Considering the broad influence of U1 snRNP on biological processes, we selected a short-term, 4-hour treatment for subsequent studies. The qRT-PCR results indicated that the lowest Cq value, corresponding to maximal blocking efficiency, was achieved with 25 μM U1 AMO (Fig. 5B). Therefore, 25 μM was selected for subsequent experiments.

**Fig. 5.**
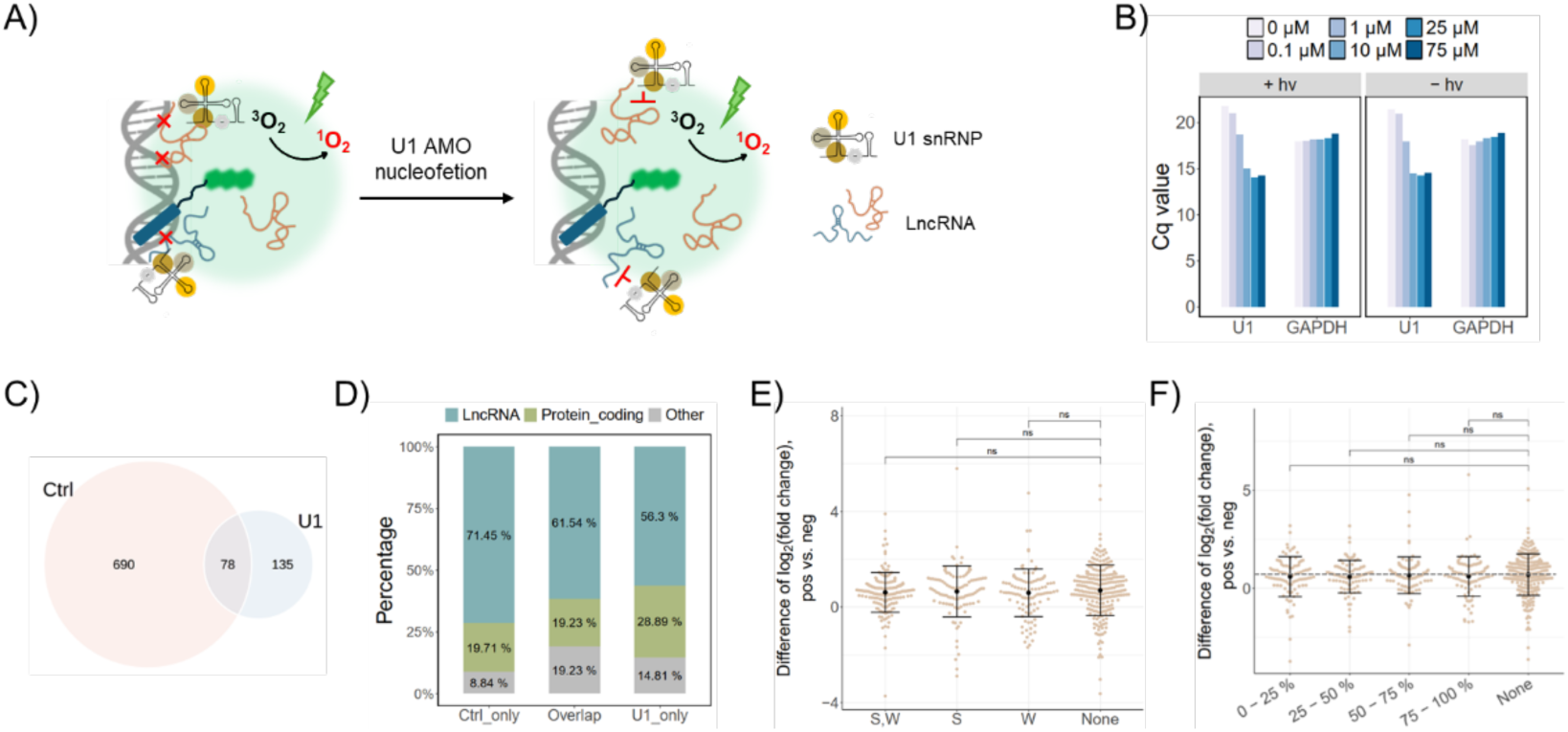
PCRD-seq characterization of the chromatin-associated transcriptome alterations resulted from U1 inhibition. (A) Schematic illustration of chromatin-associated transcriptome profiling by PCRD-seq following U1 AMO nucleofection. (B) qRT-PCR analysis following RNase H protection assay for U1 AMO concentration titration experiment. (C) Venn diagram showing the overlap of caRNAs identified following U1 AMO versus control AMO nucleofection. (D) Gene biotype compositions of caRNAs in Ctrl_only, U1_only and Overlap groups. (E) The log_2_(fold change) differences between U1 and control AMO groups for transcripts in different 5’ SS strength categories. Differences were analyzed using Wilcoxon test. (F) The log_2_(fold change) differences between U1 and control AMO groups for transcripts in different 5’ SS frequency quantiles. Differences were analyzed using Wilcoxon test.

To determine if U1 inhibition affects HoeDBF localization or labeling capability, we performed live-cell and fixed-cell confocal imaging after 4-hour nucleofection. Live-cell imaging confirmed that HoeDBF itself was specifically localized to nuclei (Fig. S5A), and fixed-cell imaging confirmed the nuclear localization of the labeled biomolecules (Fig. S5B). These results indicated that U1 inhibition did not compromise the nuclear localization or labeling specificity of HoeDBF.

We next proceeded to profile and compare the chromatin-associated transcriptomes of cells subjected to U1 or control AMO nucleofection. Outliers were excluded (Fig. S6A) for caRNA identification. The results revealed a dramatic reshuffling upon U1 inhibition. Only 213 caRNAs were detected after U1 inhibition (Fig. S6B), less than one-third of the 768 caRNAs identified in control cells (Fig. S6C), indicating global “chromatin release” of caRNAs. Venn analysis showed a modest overlap between the caRNAs identified between these two conditions both for all RNA biotypes (Fig. 5C) and for the lncRNA subset (Fig. S6D). For subsequent analysis, we categorized the caRNAs into three groups: 1) Ctrl_only: those exclusive to control AMO group, presumed to be released from chromatin due to U1 inhibition; 2) Overlap: those detected in both groups, considered unaffected by U1 inhibition; and 3) U1_only: those exclusive to the U1 inhibition group, hypothesized to be chromatin-bound as a result of U1 inhibition. Biotype composition analysis revealed a higher proportion of protein-coding RNAs in the U1_only group compared to the other two groups (Fig. 5D), possibly because U1 inhibition impairs mRNA splicing, consequently attenuating their nuclear export.

We next sought to determine whether U1-mediated lncRNA chromatin retention depends on the presence or strength of predicted 5’ splice site (5’ SS) motifs within lncRNAs, as the model suggests in mESCs ^38^. Based on established methods ^38, 40, 41^, we predicted and classified 5’ SS motifs in the identified chromatin-associated lncRNAs as strong or weak. We then categorized lncRNAs into four groups: “S” (containing strong 5’ SS only), “W” (containing weak 5’ SS only), “S,W” (containing both), or “None” (no predicted 5’ SS). We hypothesized that U1 inhibition would specifically cause “chromatin release” for lncRNAs containing 5’ SS motifs. Therefore, we hypothesized that lncRNAs without 5’ SS would show comparable log_2_(fold change) in U1 inhibition and control groups, while those harboring at least one 5’ SS motif would exhibit an increase in the log_2_(fold change) following U1 inhibition. However, contrary to our expectation, the results revealed that U1 inhibition also led to an increase in the log_2_(fold change) of caRNAs lacking a 5’ SS, i.e., the difference of the log_2_(fold change) between U1 inhibition and control groups is greater than 0 (Fig. 5E). Moreover, the magnitude of this increase was comparable across all four categories (Fig. 5E). We next analyzed the 5’ SS frequency of the chromatin-associated lncRNAs, where the lncRNAs were divided into quartiles according to their 5’ SS frequency per kilobase. Consistent with the results of 5’ SS strength analysis, the log_2_(fold change) differences between U1 inhibition and control groups were comparable across lncRNAs in all quantiles and the “None” group (Fig. 5F).

To validate the results of our 5’ SS-related analysis, we analyzed the RNA-seq data of the mESCs subjected to U1 or control AMO nucleofection ^38^. Consistent with our own findings, no biologically relevant changes in the chromatin enrichment factor were detected across lncRNAs in different 5’ SS frequency quantiles or in the “None” group following U1 inhibition (Fig. S6E). We speculated that this is due to the lack of a direct causal relationship between U1 binding and the presence of a predicted 5’ SS in a transcript. The 5’ SS recognition by U1 snRNA depends on the base-pairing between the 5’-end sequence of U1 and the 5’ SS sequence of target transcripts. However, the base-pairing between them is highly diverse. In addition to the canonical base-paring register, alternative ones, such as shifted ^42, 43^ and bulged ^43, 44^ register, have also been reported. Moreover, the 5’ SS may be sequestered by the secondary structures of its host transcript, thus preventing access by U1 snRNA ^43^. Therefore, a computationally predicted weak 5’ SS does not necessarily preclude efficient recognition and binding by U1 snRNP, and vice versa.

Overall, our PCRD-seq identified a set of caRNAs whose chromatin retention depends on U1 snRNP. However, the chromatin retention of a given lncRNA does not depend on the presence, strength, or frequency of any predicted 5’SS within its sequence, likely due to the known complexity of U1 snRNP recognition involving non-canonical and structure-sensitive interactions.

### Characterizing chromatin-associated transcriptomes of two ovarian cancer cell lines using PCRD-seq

Given the regulatory potential of caRNAs, we hypothesized that their differential chromatin association could contribute to cancer phenotypes. Ovarian cancer has the highest case-fatality rate among gynecologic malignancies, owing primarily to the fact that most patients are diagnosed at an advanced stage with extensive peritoneal metastasis ^45^. Wong’s team derived a pair of single-cell clones with opposite metastatic phenotypes—highly metastatic (HM) and non-metastatic (NM)—from the human ovarian cancer cell line HeyA8 ^46^. To investigate the potential role of RNA chromatin localization in conferring their distinct metastatic capability, we profiled the chromatin-associated transcriptome of the two cell lines using our PCRD-seq.

We first confirmed the correct spatial distribution of HoeDBF molecules and labeled biomolecules in both lines by live-cell (Fig. S7A) and fixed-cell (Figs. S7B and S7C) confocal imaging. Next, qRT-PCR following HoeDBF labeling was conducted to test the crosslinking-induced caRNA depletion effect in these cell lines. Similar to the observations in HEK293T cells, increases in the Cq values of chromatin—but not cytosolic or mitochondrial—marker genes were observed following HoeDBF labeling, confirming the depletion effect in both cell lines (Fig. 6A).

**Fig. 6.**
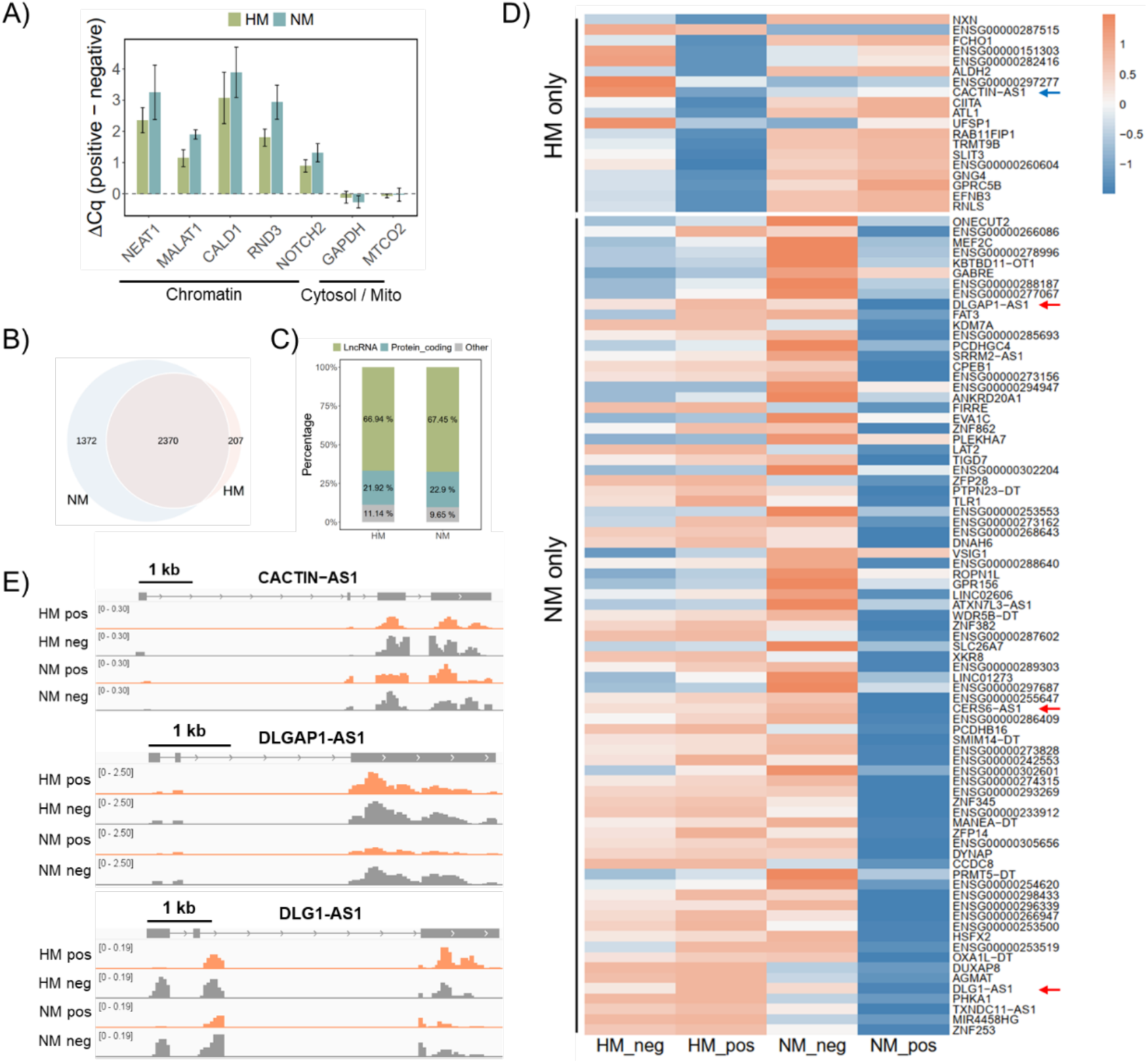
Profiling of caRNAs in HM and NM cells using PCRD-seq. (A) qRT-PCR results showing the crosslinking-induced Cq value change following HoeDBF labeling in HM and NM cells. (B) Venn diagram showing the overlap between caRNAs identified in HM and NM cells. (C) Gene biotype compositions of caRNAs identified in HM and NM cells. (D) Heatmap showing the expression patterns of caRNAs within HM_only (upper panel) and NM_only (lower panel) categories. (E) Genome tracks of three representative caRNAs indicated by blue and red arrows in Fig. D.

Subsequently, transcriptome-wide caRNA profiling was conducted for the two cell lines (Fig. S8A). Applying the criteria of log_2_(fold change) ≤ -1 and FDR < 0.05, 2,577 and 3,742 caRNAs were identified in HM (Fig. S8B) and NM (Fig. S8C) cells, respectively. The Venn diagram revealed a substantial overlap between the two sets of caRNAs (Fig. 6B). The biotype composition analysis indicated that both caRNA sets are composed predominantly of lncRNAs (Fig. 6C). We then defined cell line-exclusive caRNAs (“HM_only” and “NM_only”) based on specific criteria. HM_only caRNAs were defined as: 1) the log_2_(fold change) ≤ -1 and FDR < 0.05 in HM cells; 2) the log_2_(fold change) ≥ 0 in NM cells. Conversely, NM_only caRNAs were defined as: 1) the log_2_(fold change) ≤ -1 and FDR < 0.05 in NM cells; 2) the log_2_(fold change) ≥ 0 in HM cells. A heatmap illustrates the relative abundance of these cell line-exclusive caRNAs within labeled and control groups, reflecting their distinct chromatin association patterns (Fig. 6D). Three representative examples are shown in genome tracks (Fig. 6E).

Notably, several NM_only caRNAs were implicated in oncogenesis or metastasis. The lncRNA DLGAP1-AS1 (indicated by red arrow in Fig. 6D) was reported to be a malignancy promoter in various cancers, including gastric cancer ^47^, glioblastoma ^48^ and hepatocellular carcinoma ^49^, where it enhances tumorigenic properties such as cell proliferation, migration or invasion. Its primary oncogenic mechanism involves acting as a molecular sponge, whereby DLGAP1-AS1 competitively binding with specific microRNAs (miRNAs), thus altering the expression of their mRNA targets and modulating the downstream cancer-related signaling pathways ^47–49^. This mechanism is also employed by DLG1-AS1 and CERS6-AS1 (indicated by red arrows in Fig. 6D) to drive the tumorigenesis and progression of various cancers ^50–53^. Since the mRNA targets of the miRNAs are primarily localized in cytoplasm, we speculated that the interaction between these lncRNAs and their miRNA targets should also occur in cytoplasm. However, the heatmap (Fig. 6D) and genome tracks (Fig. 6E) revealed that although these lncRNAs exhibit comparable total RNA expression level between HM and NM cells (comparable abundance in HM_neg and NM_neg groups), they were restricted at chromatin in NM cells (significantly lower abundance in NM_pos group compared to NM_neg group). We therefore hypothesize that these lncRNAs promote metastasis in HM cells. In contrast, in NM cells, their chromatin localization prevents them from interacting with their miRNA targets, thereby suppressing their malignancy promoter function.

In summary, we profiled the chromatin-associated transcriptomes of two ovarian cancer cell lines with opposite metastatic capability using our PCRD-seq. Several RNAs involved in oncogenesis or metastasis were found to exhibit distinct chromatin localization patterns between the two cell lines, suggesting their potential roles in contributing to the observed differences in metastatic capability.

## Discussion

In this study, we developed PCRD-seq, an enrichment-free method for profiling chromatin-associated RNAs. By leveraging the proximity crosslinking induced by singlet oxygen (^1^O_2_) during HoeDBF-mediated labeling, PCRD-seq enables the identification of caRNAs based on their specific depletion from the soluble RNA pool upon TRIzol extraction. This streamlined approach overcomes several limitations of existing techniques, including complex protocols, antibody dependency, and high-input requirements, offering a simpler and more accessible tool for studying chromatin-associated RNAs (caRNAs).

As with traditional proximity labelling approaches, PCRD-seq captures caRNAs under near-native conditions in living cells, avoiding the potential artifacts associated with aldehyde fixation used in many immunoprecipitation- or proximity ligation-based methods. However, unlike traditional proximity labeling methods, which rely on a cumbersome, multi-step enrichment-based protocol, PCRD-seq identifies caRNAs by directly detecting crosslinked RNAs through their specific depletion, significantly simplifying sample handling procedure and thereby minimizing technical variability. More importantly, PCRD-seq circumvents the high-input requirement of the enrichment protocol in traditional proximity labelling technique. Its low-input compatibility positions it well for applications in primary cells or limited clinical samples. Our comparative analyses indicate that PCRD-seq preferentially identifies RNAs with stronger chromatin-association strength (Figs. 3G and 3H), suggesting it is particularly suited for studying tightly chromatin-tethered transcripts. This property, combined with its simplicity, make PCRD-seq a valuable addition to the methodological toolkit for caRNA biology.

Beyond general caRNA profiling, we demonstrated the utility of PCRD-seq in addressing specific biological questions. By applying it to the caRNA profiling in U1 snRNA inhibited HEK293T cells, we confirmed the role of U1 snRNP in global chromatin retention of lncRNAs in human cells, extending previous findings in mouse systems. Notably, our data suggest that this retention mechanism is not strictly predictable by the presence, strength or frequency of canonical 5’ splice site motifs within the target RNAs (Figs. 5E and 5F). This observation aligns with the known complexity of U1 snRNP recognition, which involves non-canonical and structure-sensitive interactions, and underscores the value of experimental profiling over purely sequence-based predictions.

In applying PCRD-seq to a matched pair of ovarian cancer cell lines with opposing metastatic capabilities, we identified distinct chromatin association patterns for several oncogenic lncRNAs (Figs. 6D and 6E). The specific chromatin retention of transcripts such as DLGAP1-AS1, DLG1-AS1 and CERS6-AS1 in non-metastatic cells suggests a model whereby spatial sequestration at chromatin may limit their cytoplasmic availability to act as miRNA sponges, thereby suppressing their pro-metastatic functions. These findings highlight how differential subnuclear localization of regulatory RNAs could contribute to phenotypic variation in cancer, a dimension that is often overlooked in bulk transcriptomic studies.

The adaptability of PCRD-seq was evidenced by its successful application to profile RNAs associated with nuclear lamina, using a HaloTag-targeted system. This result suggests the potential of PCRD-seq in profiling RNAs associated with other chromatin subdomains. However, our data also indicate that the efficiency of proximity crosslinking-induced depletion is highly context-dependent. It is most pronounced in DNA-rich environments including chromatin and mitochondria, where DNA may act as a scaffold for the formation of large DNA-protein-RNA crosslinking complex. This property is a strength for studying chromatin- and mitochondria-localized transcripts with high specificity but may limit the method’s sensitivity for profiling RNAs in other subcellular regions.

Looking forward, PCRD-seq can be readily adapted to target other chromatin subdomains—such as specific histone modifications, transcription factor binding sites, or nuclear bodies—by recruiting the photosensitizer DBF to the relevant protein tags. Proximity crosslinking-induced depletion effect can potentially be extended to chromatin-associated protein profiling, which can be further integrated with PCRD-seq to facilitate the analysis of RNA-protein interactions at chromatin. The application in addition to DNA-rich subcellular locations can be expanded by the development of an alternative specific depletion-based approach, which integrates our ^1^O_2_-mediated proximity labelling with the meCLICK-Seq described in the introduction.

In conclusion, PCRD-seq provides a simple, versatile, and enrichment-free strategy for profiling caRNAs. By leveraging the detection of specific RNA depletion to overcome enrichment challenges, this approach facilitates the study of RNA chromatin localization patterns. It thus unlocks new possibilities for investigating how caRNAs function in both normal physiology and disease states.

## Notes

### Competing Interest Statement

The authors have declared no competing interest.

### Funding

- the Hong Kong Research Grants Council: 17318422, 17308925
- the Laboratory for Synthetic Chemistry and Chemical Biology under the Health@- InnoHK Program launched by the Innovation and Technology Commission

