## supporting information for "PCRD-seq: Proximity Crosslinking-induced RNA Depletion for Low-Input Subcellular Transcriptome Profiling"

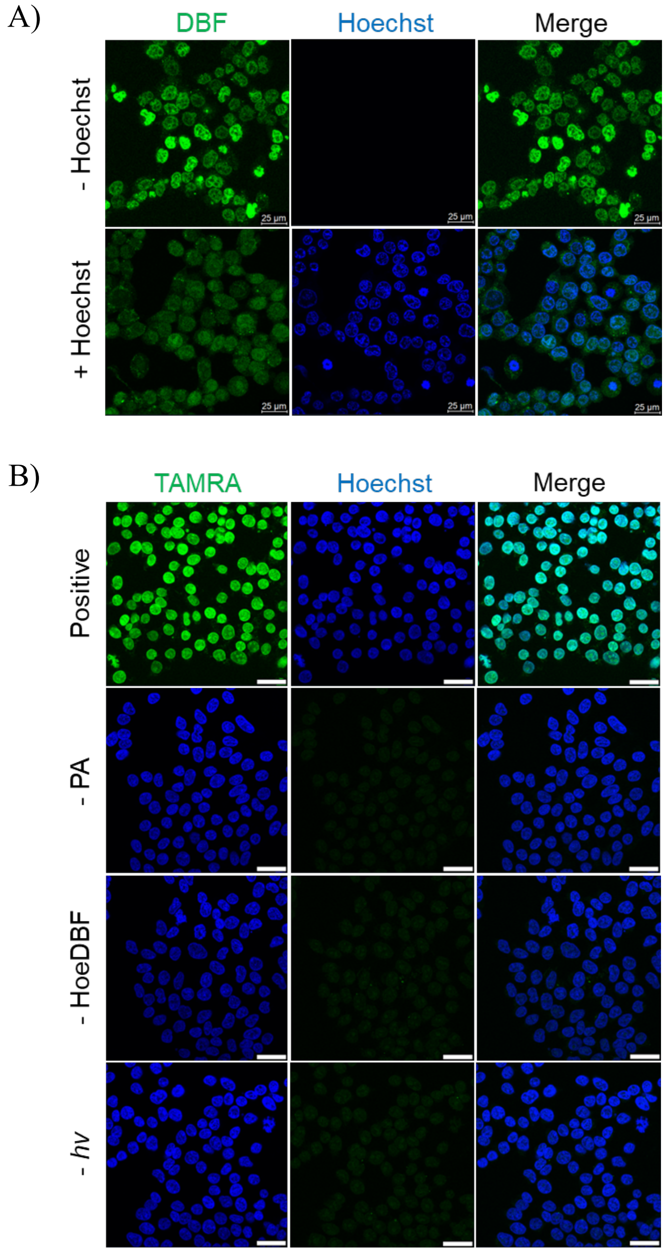


**Fig. S1** (A) Fluorescence confocal imaging of living cells after incubation with HoeDBF. Counterstaining with Hoechst 33342 was performed where indicated. The green signal represents HoeDBF molecules bound to chromatin. The blue signal represents Hoechst 33342 nucleic acid stain. Scale bar: 25 µm. (B) Fluorescence confocal imaging in fixed cells after HoeDBF labeling. TAMRA was ligated to labeled biomolecules for visualization. Hoechst 33342 was used for the visualization of nucleus. "Positive" indicates cells subjected to HoeDBF labeling, while "- PA", "- HoeDBF" and "- *hv*" indicate the respective negative controls. Scale bar: 25 µm.

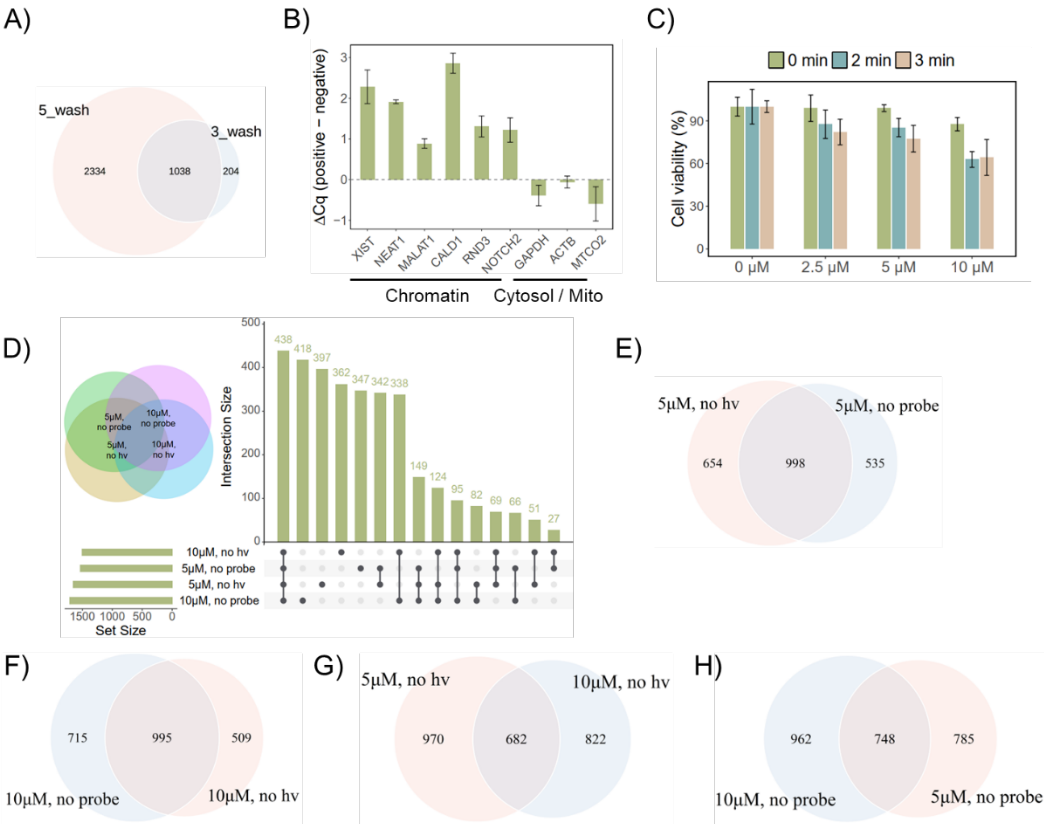


**Fig. S2** (A) Venn diagram illustrating the overlap of caRNAs captured under different washing stringency conditions. "3_wash" and "5_wash" refer to experiments where beads bound to biotinylated RNAs were subjected to three and five wash cycles, respectively. (B) qRT-PCR analysis of RNAs isolated from HoeDBF labeled cells (positive) and negative control (negative). (C) Cell viability test using CCK-8 assay under different labeling conditions. (D) The UpSet plot indicates the number of caRNAs identified under each labeling condition and their intersections. The inset Venn diagram provides a general overview of the overlaps between the conditions. (E-H) Venn diagram showing the overlap of caRNAs between each two labeling conditions.

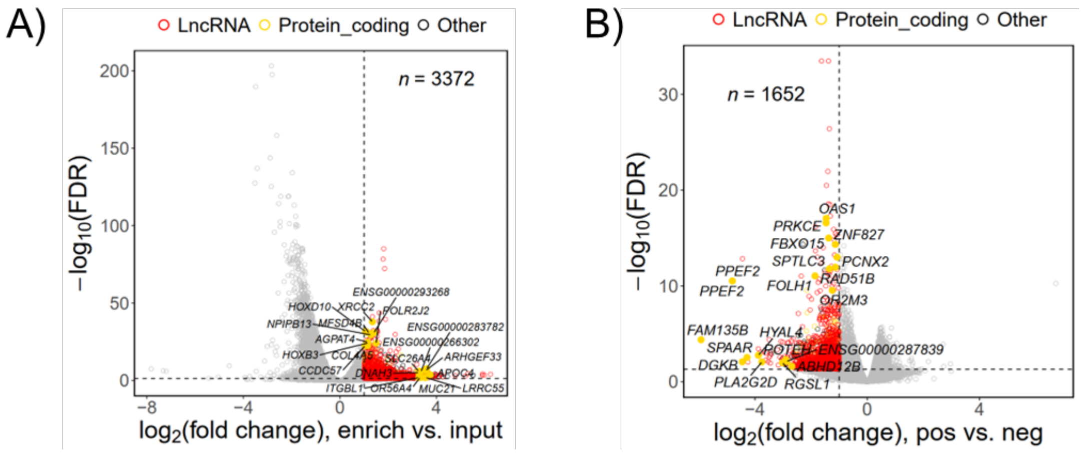


**Fig. S3** (A) Volcano plot showing the differential expression between enriched and input samples for HoeDBF-mediated enrichment-based method. Red and yellow dots indicate significantly up-regulated lncRNAs and protein-coding RNAs ($\log_2$(fold change) $\geq$ 1, FDR $<$ 0.05) respectively, i.e., caRNAs. The yellow solid dots highlight the top 10 protein-coding RNAs with the lowest FDR or highest $\log_2$(fold change) values. Most of the highlighted RNAs are not linked to lamina. (B) Volcano plot showing the differential expression between labeled (positive) and control (negative) groups for HoeDBF-mediated depletion-based method. Red and yellow dots indicate significantly down-regulated lncRNAs and protein-coding RNAs ($\log_2$(fold change) $\leq$ -1, FDR $<$ 0.05) respectively, i.e., caRNAs. The yellow solid dots highlight the top 10 protein-coding RNAs with the lowest FDR or $\log_2$(fold change) values. Most of the highlighted RNAs are not linked to lamina.

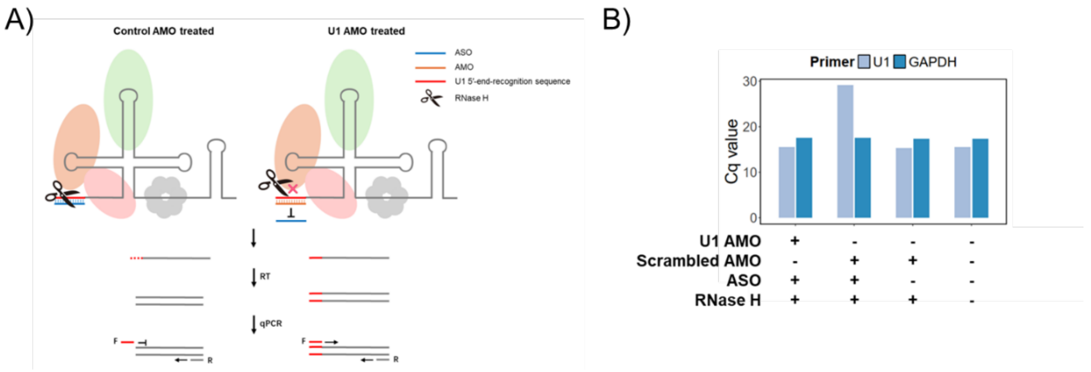


**Fig. S4** (A) The principle of the method used for U1 recognition sequence blocking efficiency assessment which integrates RNase H protection assay and qRT-PCR analysis. Unlike natural antisense oligonucleotides (ASO), an AMO is an unnatural nucleic acid analog. Consequently, while an RNA-DNA duplex is a substrate for RNase H digestion, an RNA-AMO duplex is not. We therefore reasoned that the U1 5'-end recognition site would remain intact in U1 AMO-treated samples (as it is protected by the AMO) but would be digested in control AMO-treated samples upon addition of RNase H and ASO. For the subsequent qRT-PCR analysis, a pair of primers targeting U1 snRNA was designed, among which the forward primer target to U1 5'-end recognition sequence. Therefore, in control AMO-treated samples where this recognition site is digested, the target sequence cannot be amplified, resulting in a significantly higher Cq value compared to the U1 AMO-protected samples. (B) The qRT-PCR analysis following RNase H protection assay using total RNAs. The results showed that only in samples pre-treated with U1 AMO (and not in samples treated with control AMO) was the Cq value maintained upon addition of RNase H and ASO, comparable to the baseline level in samples without RNase H and ASO, thereby validating this method.

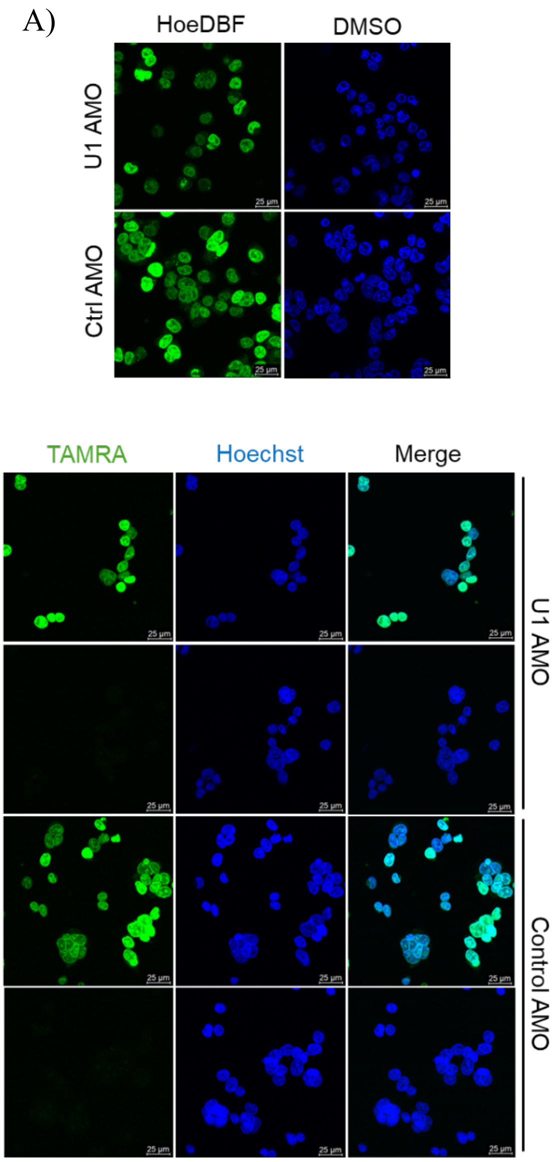


**Fig. S5** (A) Fluorescence confocal imaging in living cells 4 hours after U1 or control AMO nucleofection. Green signal represents HoeDBF molecules binding to chromatin. Blue signal represents Hoechst 33342 nucleic acid stain. Scale bar: 25 μm. (B) Fluorescence confocal imaging in fixed cells 4 hours after U1 or control AMO nucleofection. TAMRA was ligated to labeled biomolecules for visualization. “Positive” indicates cells subjected to HoeDBF labeling, while “- PA” indicates negative control. Scale bar: 25 μm.

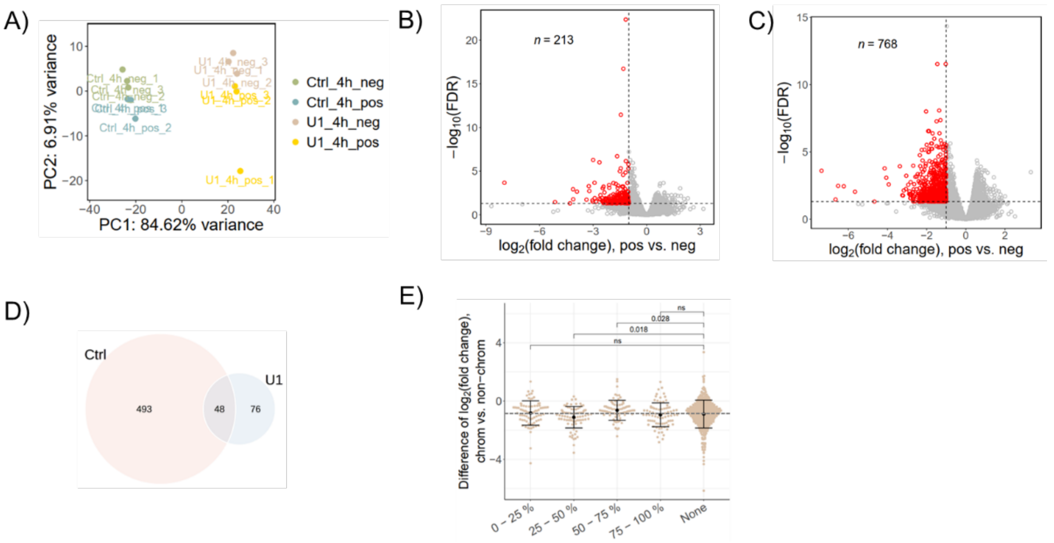


**Fig. S6** (A) PCA based on normalized read counts for samples subjected to U1 or control AMO nucleofection. Sample nomenclature indicates the AMO treatment and labeling status: "U1_4h_pos" and "Ctrl_4h_pos" refer to samples nucleofected with U1 AMO or control AMO, respectively, followed by HoeDBF labeling. Conversely, "U1_4h_neg" and "Ctrl_4h_neg" denote corresponding samples that did not undergo HoeDBF labeling. One replicate from the U1_4h_pos group (U1_4h_pos_1) was identified as an outlier and excluded, leaving only two replicates. To maintain parity in replicate numbers for a fair statistical comparison, one replicate from Ctrl_4h_pos group (Ctrl_4h_pos_1) was accordingly also excluded. (B and C) Volcano plots showing the differentially expressed genes (DEGs) between HoeDBF labeled (positive) and control (negative) groups following U1 AMO (B) or control AMO (C) nucleofection. Those defined as caRNAs ($\log_2$(fold change) $\leq$ -1, FDR $<$ 0.05) are highlighted in red. (D) Venn diagram showing the overlap between chromatin-associated lncRNAs identified in samples subjected to U1 AMO and control AMO nucleofection. (E) The $\log_2$(fold change) differences between U1 and control AMO nucleofected mESCs for transcripts in different 5' SS frequency quantiles. Consistent with the findings in our own data, we observed a decline in the chromatin enrichment factor for chromatin-associated lncRNAs both with and without a 5' SS. Although U1 inhibition caused statistically significant differences in the chromatin enrichment factor for lncRNA in some quantiles (e.g., 25–50% and 50–75%) compared to those without a 5’ SS, the directions of the changes were inconsistent and, in some cases (e.g., 50–75%), opposite to expectations. Furthermore, the magnitude of these alterations was minimal. Therefore, we speculated that this change is driven by random variation rather than by U1 inhibition.

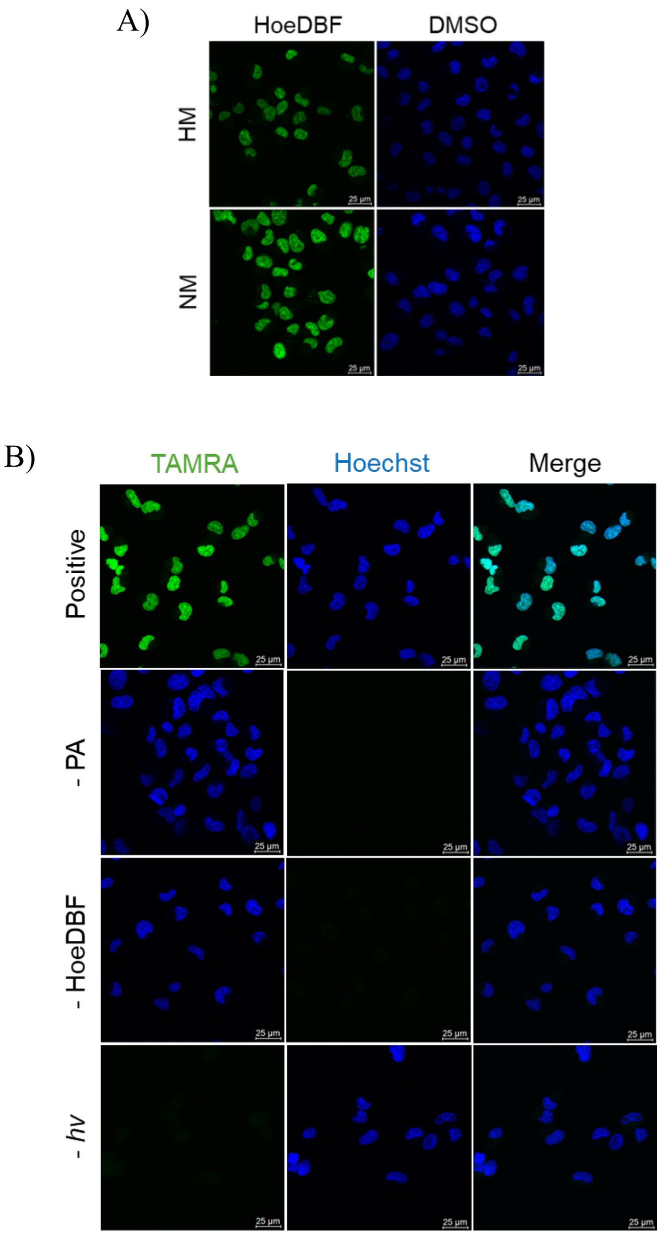

C)

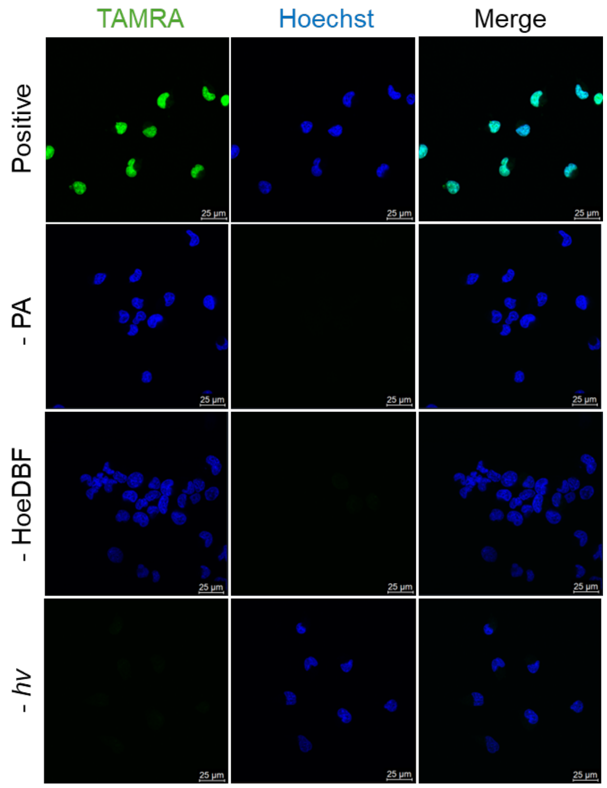


**Fig. S7** (A) Fluorescence confocal imaging in living HM and NM cells after incubation with HoeDBF. Green signal represents HoeDBF molecules bound to chromatin. Blue signals represent Hoechst 33342 nucleic acid stain. Scale bar: 25 μm. (B and C) Fluorescence confocal imaging in fixed HM (B) and NM (C) cells after HoeDBF labeling. TAMRA was ligated to labeled biomolecules for visualization. Hoechst 33342 was used for the visualization of nucleus. "Positive" indicates cells subjected to HoeDBF labeling, while "- PA", "- HoeDBF" and "- *hv*" indicate the respective negative controls. Scale bar: 25 μm.

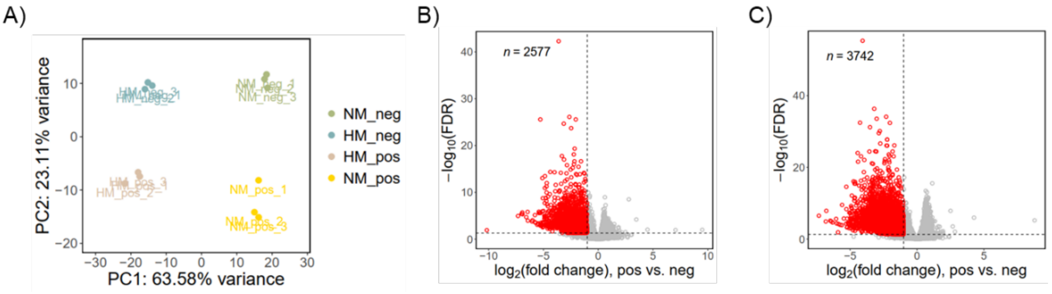


**Fig. S8** (A) PCA based on normalized read counts. (B and C) Volcano plots showing DEGs between labeled (positive) and control (negative) samples in HM (B) and NM (C) cells, respectively. Red dots indicate the transcripts defined as caRNAs ($\log_2$(fold change) $\leq$ -1, FDR $<$ 0.05).

**Table S1** Descriptions for different HoeDBF labeling conditions

| Labeling condition | Abbreviation |
|---|---|
| Incubation with 5 μM HoeDBF followed by 2 min green light irradiation versus incubation with 5 μM HoeDBF without light irradiation. | 5 μM, no *hv* |
| Incubation with 5 μM HoeDBF followed by 2 min green light irradiation versus incubation with DMSO followed by 2 min green light irradiation. | 5 μM, no HoeDBF |
| Incubation with 10 μM HoeDBF followed by 3 min green light irradiation versus incubation with 10 μM HoeDBF without light irradiation. | 10 μM, no *hv* |
| Incubation with 10 μM HoeDBF followed by 3 min green light irradiation versus incubation with DMSO followed by 3 min green light irradiation. | 10 μM, no HoeDBF |

### MATERIALS AND METHODS

#### Cell culture

HEK293T and U2OS cells were purchased from American Type Culture Collection (ATCC). HEK293T cells were cultured in Dulbecco's Modified Eagle Medium (DMEM; Gibco, 11965092) supplemented with 10% fetal bovine serum (FBS; Gibco,

10270106) and 1% penicillin/streptomycin (P/S; Gibco, 15140122) at 37 ºC under 5% $CO_2$. U2OS cells were cultured in McCoy's 5A (Modified) Medium (McCoy's 5A; Gibco, 16600082) supplemented with 10% FBS and 1% P/S at 37 ºC under 5% $CO_2$. HM and NM cells were a generous gift from Professor Alice Sze Tsai WONG (Affiliation, The University of Hong Kong). These two cell lines were cultured in RPMI 1640 medium (Gibco, 11875093) supplemented with 5% FBS and 1% P/S at 37 ºC under 5% $CO_2$. The homemade Flp-In-HEK293/HaloTag7-LMNA and Cre-loxP-HeLa/FAP-G3BP1 stable cell lines were maintained in their respective selection media at 37°C under 5% $CO_2$. The Flp-In-HEK293/HaloTag7-LMNA cell line was cultured in DMEM with 10% FBS, 1% P/S, 10 μg/mL blasticidin S, and 200 μg/mL hygromycin B, while the Cre-loxP-HeLa/FAP-G3BP1 cell line was cultured in DMEM with 10% FBS, 1% P/S, and 5 μg/mL puromycin.

**Stable Flp-In-HEK293/HaloTag7-LMNA cell line generation and transgene expression induction**

HEK293 cell line stably expressing Halo-LMNA was generated using Flp-In system. The Flp-In-HEK293 cell line, kindly provided by Professor Gary Ying Wai Chan (Affiliation, The University of Hong Kong), served as the host for generating the inducible stable cell line. The pcDNA5/FRT/TO-HaloTag7-LMNA plasmid was constructed by Hunan Fenghui Biotechnology Co., Ltd. For transfection, the Flp-In-HEK293 cells were seeded in a 6-well plate and cultured in standard culture medium (DMEM supplemented with 10% FBS and 1% P/S) further supplemented with 100 μg/mL zeocin and 15 μg/mL blasticidin S at 37°C under 5% $CO_2$. Upon reaching 50–60% confluence, the cells were co-transfected with 1.8 μg pOG44 plasmid and 200 ng pcDNA5/FRT/TO-HaloTag7-LMNA plasmid using Lipofectamine 2000 Reagent (Invitrogen, 11668019), following the manufacturer's instructions. At 48 hours post-transfection, the culture was assessed for density. If the cells were below 25% confluence, the existing medium was replaced with the selection medium (standard culture medium supplemented with 10 μg/mL blasticidin S, and 200 μg/mL hygromycin B), and maintained in the same 6-well plate. If the confluence exceeded 25%, the cells were trypsinized, subcultured into a 6-cm dish, and maintained in the same selection medium. The selection medium was refreshed every 2-3 days until cells reached 90% confluence. The resulting polyclonal stable cells were then cryopreserved for long-term storage or subcultured for subsequent experimental use.

The homemade Flp-In-HEK293/HaloTag7-LMNA stable cell line was maintained in the selection medium. To induce transgene expression, the existing medium was replaced with induction medium (selection medium supplemented with 2 μg/mL doxycycline) 24 hours prior to experimentation.

**Stable Cre-loxP-Hela/FAP-G3BP1 cell line generation and transgene expression induction**

The Hela cell line stably expressing FAP-G3BP1 fusion protein was established using Cre-loxP recombination system. The host HeLa cells, containing the genomic loxP cassette, were seeded in a 6-well plate and cultured in standard culture medium (DMEM supplemented with 10% FBS and 1% P/S). At 70–80% confluence, the cells were co-transfected using Lipofectamine 2000 Reagent (Invitrogen, 11668019) with two plasmids: the FAP-G3BP1 donor plasmid (constructed by Hunan Fenghui Biotechnology Co., Ltd) and the pBT140 plasmid, following the manufacturer's protocol. Briefly, 6 μL of Lipofectamine 2000 reagent was diluted in 150 μL of Opti-MEM medium (Gibco, 31985070). Separately, a DNA mixture containing 2 μg of the FAP-G3BP1 plasmid and 100 ng of the pBT140 plasmid was diluted in another 150 μL of Opti-MEM. Next, 150 μL of diluted DNA and 150 μL of diluted Lipofectamine 2000 Reagent were mixed, and incubated for 5 min at room temperature. Subsequently, 250 μL of the mixture was added to each well. Following 24-hour incubation, the medium was replaced by fresh culture medium and cells were incubated for another 24 hours. Selection was then initiated by replacing the medium with a selection medium (standard culture medium supplemented with 2.5 μg/mL puromycin) and incubating for 48 hours, followed by maintenance in a higher concentration of puromycin (5 μg/mL) for 12 days. Cells were passaged into 6-cm dishes as needed during the selection period. The resulting polyclonal stable cells were then cryopreserved for long-term storage or subcultured for subsequent experimental use.

The established Cre-loxP-Hela/FAP-G3BP1 cells were cultured in selection medium containing 5 μg/mL puromycin. To induce the expression of FAP-G3BP1 fusion protein, the culture medium was replaced with induction medium (standard culture medium supplemented with 5 μg/mL puromycin and 1 μg/mL doxycycline) 48 hours prior to experimentation.

**HoeDBF labeling in living cells for HoeDBF-mediated enrichment-based**

**experiments**

Cells were cultured in Poly-D-Lysine (PDL; Millipore, A-003-E)-treated dishes as described above and subjected to HoeDBF labeling upon reaching 70–80% confluence. The culture medium was removed, and the cells were washed once with Dulbecco's phosphate-buffered saline (DPBS; Gibco, 14190144). Subsequently, cells were incubated at 37 °C for 30 min with a solution containing 5 μM HoeDBF, prepared in Hanks' Balanced Salt Solution (HBSS; Gibco, 14175095) with 1% DMSO. Control cells were incubated under identical conditions with HBSS containing 1% DMSO alone. Cells were then washed once with DPBS to remove residual HoeDBF, followed by two 20-minute incubations in culture medium at 37 °C to facilitate the clearance of unbound HoeDBF molecules in cells. Cells were then washed once again with DPBS and incubated with 3 mM PA in HBSS for 2 min. Finally, while still immersed in the PA solution, the cells were exposed to green light (520 nm, 150 $W/m^2$) for 2 min. The labeled cells were used for subsequent experiments.

**HoeDBF labeling in living cells for HoeDBF-mediated depletion-based experiments**

Cells were cultured in PDL-treated dishes as described above and subjected to HoeDBF labeling upon reaching 70–80% confluence. The culture medium was removed and the cells were washed once with DPBS. Subsequently, cells were incubated at 37 °C for 30 min with a solution containing 5 μM or 10 μM HoeDBF, prepared in HBSS with 1% DMSO. Cells were then washed once with DPBS to remove residual HoeDBF, followed by two 20-minute incubations in culture medium at 37 °C to facilitate the clearance of unbound HoeDBF molecules from cells. After an additional DPBS wash, the cells, while immersed in HBSS, were exposed to green light (520 nm, 150 $W/m^2$) for 2 or 3 min by sandwiching them between two LED light sources. Under the respective experimental settings, control cells received the following treatments: 1) no probe: incubation with HBSS containing 1% DMSO alone instead of HoeDBF under identical conditions; 2) no *hv*: no exposure to green light. The labeled cells were then used for subsequent analysis.

**Nuclear lamina-associated RNA labeling for PCRD-seq**

The homemade Flp-In-HEK293/HaloTag7-LMNA cells were seeded in 6-cm dishes pre-treated with PDL and cultured in selection medium (DMEM supplemented with 10%

FBS, 1% P/S, 10 μg/mL blasticidin S, and 200 μg/mL hygromycin B) at 37°C under 5% $CO_2$. Prior to labeling, cells were induced with induction medium (selection medium supplemented with 2 μg/mL doxycycline) for 24 hours to express the HaloTag7-LMNA fusion protein. Upon reaching 80% confluence, the cells were processed for labeling. Briefly, the cells were first washed once with HBSS and then incubated with 2 μM Halo-DBF in HBSS for 15 min at 37°C. Following a quick wash with HBSS, cells were subjected to two 20-minute incubations in standard culture medium (without blasticidin S and hygromycin B). Cells were then washed once with HBSS and exposed to green light (520 nm, 150W/$m^2$) for 1 min by sandwiching them between two LED light sources, while control samples were not. Cells were then lysed in TRIzol (Invitrogen, 15596018) and stored at - 80°C after two final HBSS washes.

**Mitochondrion-, endomembrane- and lysosome-associated RNA labeling for PCRD-seq**

U2OS cells were seeded in 6-cm dishes and cultured to 90% confluence in McCoy's 5A medium supplemented with 10% FBS and 1% P/S at 37°C under 5% $CO_2$. To label organelle-specific RNA, the medium was replaced with fresh medium containing the respective probes: 0.1 μM SeNB-Amide (for endomembrane), 2.5 μM SeNB-Mito (for mitochondria), and 0.1 μM SeNB-Lyso (for lysosomes). Following 30-min incubation, cells were washed with HBSS and irradiated with 730 nm light (~ 400 W/$m^2$) in McCoy's 5A medium without FBS or P/S. Control groups were maintained in the dark. Cells were then lysed in RNAiso Plus (Takara, 9109) and immediately stored at - 80°C after two final HBSS washes.

**Stress granule-associated RNA labeling for PCRD-seq**

The homemade Cre-loxP-HeLa/FAP-G3BP1 cells were seeded in 6-cm dishes and maintained in puromycin selection medium (DMEM supplemented with 10% FBS, 1% P/S, and 5 μg/mL puromycin) at 37°C under 5% $CO_2$. To induce FAP-G3BP1 expression, cells were incubated with induction medium (DMEM supplemented with 10% FBS, 1% P/S and 1 μg/mL doxycycline) for 48 hours. Prior to labeling, cells were washed with HBSS and then incubated with 0.5 μM MGHI probe in DMEM without FBS or P/S for 20 min at 37°C. To induce stress granule assembly, the "+ stress" group was treated with 0.5 mM sodium arsenite in standard culture medium for 40 min, while the "- stress" control was incubated in culture medium. After a quick HBSS wash, the cells were irradiated with 660 nm light (~ 350 W/m$^2$) for 1 min in HBSS. Control samples were maintained in the dark. Finally, following two washes with HBSS, all samples were lysed in RNAiso Plus and immediately stored at - 80°C.

**Live-cell fluorescence confocal imaging**

Cells were cultured in 35 mm confocal dishes at 37 ºC under 5% $CO_2$ and labeled with HoeDBF at 70–80% confluence. The confocal dishes were pre-treated with PDL for HEK293T cells, whereas not for HM and NM cells. Cells were first washed once with DPBS and incubated at 37 °C for 30 min with a solution containing 5 μM HoeDBF, prepared in HBSS with 1% DMSO. Cells that incubated with HBSS containing 1% DMSO alone under the same condition served as negative control. Following a wash with DPBS, cells were subjected to two sequential 20-minute incubations in culture medium at 37 °C. Cells were then washed once with DPBS. Finally, control cells were stained with Hoechst 33342 (1:1000 dilution; Thermo Fisher Scientific) in HBSS for 10 min, whereas this step was omitted for positive samples to prevent competitive binding of Hoechst 33342 to dsDNA. The prepared samples were imaged immediately with Leica TCS SP8 confocal imaging system.

**Fixed-cell fluorescence confocal imaging**

Cells were seeded and cultured in a 6-well plate with a cover slide in the bottom of each well. The cover slides were pre-treated with PDL for HEK293T cells, whereas not for HM and NM cells. Upon reaching 70–80% confluence, cells were labeled with HoeDBF according to the labeling protocol described above for the enrichment-based method. For wild-type HEK293T cells and those subjected to U1 AMO nucleofection, negative control was prepared by omitting PA incubation. For HM and NM cells, those omitting green light irradiation were used as negative control. Following labeling, cells were washed once with DPBS and then fixed and permeabilized at room temperature for 30 min with 4% paraformaldehyde (PFA) containing 0.1% Triton X-100 in DPBS. Cells were then washed three times with DPBS and blocked with 0.1% Bovine Serum Albumin (BSA) blocking buffer at room temperature for 35 min. Following two washes with DPBS, a Copper(I)-catalyzed Azide-Alkyne Cycloaddition (CuAAC) reaction was performed to conjugate the 5-TAMRA dye to the labeled biomolecules. The reaction was performed by incubating cells in a CuAAC reaction mixture (0.5 mM $CuSO_4$, 2.5 mM THPTA, 10 mM sodium ascorbate, 10 mM TAMRA-$N_3$ in nuclease-free $H_2O$) at

37°C for 1 hour in the dark. The reagents of the CuAAC reaction solution should be added sequentially in the order listed. Cells were then washed three times with DPBS containing 0.1% Triton X-100. Finally, cells were stained with Hoechst 33342 (1:2000 dilution) for 2 min at room temperature, followed by three additional washes with DPBS. The prepared samples were mounted with VectaShield antifade mounting medium (Vector Labs, H-1000) for microscopy with Leica TCS SP8 confocal imaging system.

**RNA isolation and biotinylation though CuAAC**

HEK293T cells were labeled with HoeDBF according to the labeling protocol described above for the enrichment-based method. Total RNA was then isolated using RNAiso Plus following the manufacturer's instructions. The RNA was subsequently treated with DNase (Invitrogen, AM2239) and Proteinase K (Invitrogen, 25530049) to remove residual genomic DNA and protein. RNA biotinylation was then performed via CuAAC reaction. The reaction mixture contains 0.1 mM $CuSO_4$, 2 mM THPTA, 10 mM sodium ascorbate, 10 mM Tris PH7.0, 10 μg RNA and 2 mM biotin picolyl azide. The reaction mixture was made up to 50 μL using nuclease-free $H_2O$. The reagents were added sequentially in the order listed. The reaction mixture was incubated at 25 °C for 10 min with shaking at 500 rpm.

**RNA dot blot analysis**

A total of 300 ng biotinylated RNA was loaded onto equilibrated and dried Amersham Hybond-N+ membrane and crosslinked to the membrane using an ultraviolet crosslinker (Analytik Jena US). The membrane was then blocked using blocking buffer (0.1 g/mL sodium dodecyl sulfate (SDS), 125 mM NaCl, 17 mM $Na_2HPO_4$, and 8 mM $NaH_2PO_4$ in deionized $H_2O$) at room temperature for 30 min. Subsequently, the membrane was incubated with Streptavidin-HRP (Abcam, ab7403) in blocking buffer at room temperature for 10 min. The membrane was then washed sequentially on a shaker: twice with Wash Buffer A (blocking buffer diluted 1:10 in deionized $H_2O$) for 20 min each, followed by twice with Wash Buffer B (100 mM Tris, 100 mM NaCl, and 20 mM $MgCl_2$ in deionized $H_2O$) for 5 min each. Finally, the membrane was incubated with Clarity Western ECL Substrate (Bio-Rad, 1705060) and imaged on the ChemiDoc imaging system (Bio-Rad).

**Subcellular fractionation assay**

HEK293T cells at 70–80% confluence were first labeled with HoeDBF according to the labeling protocol for the enrichment-based method, followed by sequential extraction of cytoplasmic, nucleoplasmic, and chromatin fractions as previously described [1]. Briefly, cells were washed once with DPBS following HoeDBF labeling. Cells were then detached using 0.05% trypsin and collected after neutralization with 3 mL of culture medium. Cells were pelleted by centrifugation at 1,000 rpm for 3 min. The pellet was resuspended and gently rinsed with 1 mL of ice-cold 1x PBS. After counting, $1\times10^7$ cells were transferred to a separate 1.5 mL tube and centrifuged at 500 g for 5 min at 4°C. The supernatant was discarded, and the cells were gently resuspended in 380 μL of ice-cold Hypotonic Lysis Buffer (HLB; 10 mM Tris pH 7.5, 10 mM NaCl, 3 mM $MgCl_2$, 0.3% NP-40 (v/v), 10% glycerol (v/v) in nuclease-free $H_2O$) supplemented with 100 U of RNaseOUT (Invitrogen, 10777019). After a 10-min incubation on ice, cells were briefly vortexed and centrifuged at 1,000 g at 4°C for 3 min. The supernatant (cytoplasmic fraction) was then transferred to a new tube, with a small aliquot reserved for western blot analysis to validate fractionation purity. Following this, 1 mL of RNA precipitation solution (RPS; 150 mM sodium acetate pH 5.5 in 100% ethanol) was added to the remaining fractions immediately. The mixture was then stored at - 20°C for at least 1 hour prior to RNA isolation. The pellet was then washed three times with 1 mL of ice-cold HLB, each wash was performed by gentle pipetting followed by centrifugation at 300 g at 4°C for 2 min. After discarding the final supernatant, the pellet was resuspended in 380 μL of Modified Wuarin-Schibler buffer (MWS; 10 mM Tris-HCl pH 7.0, 4 mM EDTA, 0.3 M NaCl, 1 M urea, and 1% NP-40 (v/v) in nuclease-free $H_2O$) supplemented with 100 U of RNaseOUT. The suspension was vortexed for 30 s and incubated on ice for 5 min. Then, the suspension was subjected to an additional 30 s of vortexing and a 10-min incubation on ice. The sample was then centrifuged at 1,000 g at 4°C for 3 min. The resulting supernatant, representing the nucleoplasmic fraction, was transferred to a new tube. A small aliquot of this fraction was saved for western blot analysis. Subsequently, 1 mL of RPS was immediately added to the remaining supernatant, and the mixture was stored at - 20°C for at least 1 hour prior to RNA isolation. Finally, the pellet was washed three times with 1 mL of ice-cold MWS by vortexing for 30 s and centrifuging at 500 g at 4°C for 2 min. This final pellet constituted the pure chromatin fraction. A small portion of this fraction was saved for western blot analysis. The remaining chromatin fraction was processed directly for RNA isolation by adding 1 mL of RNAiso Plus. Total RNA isolated from the cytoplasmic, nucleoplasmic, and chromatin fractions was treated with DNase and Proteinase K, followed by biotinylation via CuAAC. The biotin signal was subsequently detected by RNA dot blot assay.

**Biotinylated RNA enrichment**

For enrichment, 80 μg of biotinylated RNA was first aliquoted into two tubes (40 μg per tube) and then processed in a 200 μL mixture for each. Specifically, Dynabeads MyOne Streptavidin C1 Beads (Invitrogen, 65002; 40 μL beads for 40 μg RNA) were pre-washed by sequential incubation with end-to-end rotation as follows: three times with 1 mL of 1× binding buffer (1 M NaCl, 0.2% Tween 20 (v/v), 100 mM Tris pH 7.5, 10 mM EDTA pH 8.0 in nuclease-free $H_2O$) for 10 min each, followed by two washes with 200 μL of Solution A (100 mM NaOH, 50 mM NaCl, 0.1% Tween 20) for 2 min each, and a final wash with 200 μL of Solution B (100 mM NaCl, 0.1% Tween 20 (v/v) in nuclease-free $H_2O$) for 2 min. The beads were finally resuspended in 2× binding buffer (2 M NaCl, 0.4% Tween 20 (v/v), 200 mM Tris pH 7.5, 20 mM EDTA pH 8.0 in nuclease-free $H_2O$). A 100 μL aliquot of the pre-washed beads in 2× binding buffer was combined with an equal volume of biotinylated RNA (40 μg) supplemented with 1 μL of RNaseOUT, thus the final 200 μL mixture was in 1× binding buffer. The mixture was incubated at room temperature for 2 h with end-to-end rotation. Subsequently, the mixture was put on a magnetic stand to separate the beads with buffer. The beads were washed three times with 1 mL of 1× binding buffer under end-to-end rotation (5 min per wash), followed by two additional washes with 1 mL of 1× binding buffer on a shaker at 50°C and 500 rpm (5 min each). To elute the enriched RNA, the beads were incubated with 50 μL of elution buffer (47.5 μL formamide, 1 μL 500 mM EDTA pH 8.0, 1.5 μL 50 mM D-biotin) at 65°C for 5 min, followed by a second incubation at 90°C for 5 min without changing the buffer. Following separation on a magnetic stand, the supernatants (containing the enriched RNA) from both tubes of the same sample were combined, yielding a 100 μL eluate. RNA was then isolated by adding 1 mL of RNAiso Plus to the eluate. The isolated RNAs were dissolved in 13.5–15.5 μL nuclease-free $H_2O$. The enriched RNA, along with input RNA (i.e., the biotinylated RNA prior to enrichment), were used for qRT-PCR and RNA-seq analysis.

**qRT-PCR analysis for enriched and input RNA**

Enriched and input RNA was first reverse transcribed using PrimeScript™ RT reagent

Kit (Takara, RR037A). Specifically, 1 μL of enriched or 300 ng of input RNA was mixed with 0.5 μL Oligo dT Primer (50 μM) and 0.5 μL Random 6 mers (100 μM). The reaction mixture was made up to 7.5 μL with nuclease-free $H_2O$ and incubated in 65°C for 5 min. Following the incubation, 2 μL of 5x PrimeScript Buffer and 0.5 μL of PrimeScript RT Enzyme Mix I were added to the reaction mixture. The resulting 10 μL mixture was then incubated at 25°C for 10 min, 42°C for 45 min, 85°C for 5 s, and finally held at 4°C. The 10 μL reaction mixture was diluted to 20–25 μL and 2 μL was used for real-time PCR analysis. Real-time PCR was performed using TB Green Premix Ex Taq (Tli RNaseH Plus) (Takara, RR420A). Specifically, 2 μL of cDNA was mixed with 6.25 μL TB Green Premix Ex Taq (2x) and primers (final concentration: 0.8 μM). The reaction mixture was made up to 12.5 μL using nuclease-free $H_2O$ and analyzed using BIORAD CFX Connect Optics Module. The fold enrichment of each target RNA was calculated as $2^{((Ct_{enrich\_DMSO} - Ct_{input\_DMSO}) - (Ct_{enrich\_HoeDBF} - Ct_{input\_HoeDBF}))}$.

**qRT-PCR analysis for the validation of crosslinking-induced caRNA depletion**

Cells were first labeled with HoeDBF according to the labeling protocol described above for the depletion-based method. For the validation in HEK293T cells, which was used to establish the PCRD-seq, the labeling conditions were 10 μM HoeDBF incubation followed by 3-min green light irradiation. For the validation in HM and NM cells, the labeling conditions were 5 μM HoeDBF incubation followed by 2-min green light irradiation. Cells without exposure to green light were used as negative control. The labeled cells were lysed immediately with RNAiso Plus for RNA isolation. The isolated total RNA was then treated with DNase and Proteinase K following the manufacturer's instructions to remove residual genomic DNA and protein. Next, 300 ng of purified total RNA was reverse transcribed using PrimeScript™ RT reagent Kit as described above. The resulting 10 μL cDNA product was then diluted to 30–40 μL with nuclease-free $H_2O$, and 2 μL of this dilution was used for qRT-PCR analysis of each target gene with TB Green Premix Ex Taq (Tli RNaseH Plus) as described above. The crosslinking-induced caRNA depletion effect was quantified by the ΔCq value, which was calculated as the difference in Cq between labeled and the control samples.

**DNase and Proteinase K treatment of cell lysate, RNA isolation and qRT-PCR analysis**

HEK293T cells were seeded and cultured in 6-well plate coated with PDL. Upon reaching 70–80% confluence, cells were labeled with HoeDBF according to the labeling protocol for the depletion-based method (labeling conditions: 10 μM HoeDBF incubation, 3-min green light irradiation). Cells without exposure to green light were used as unlabeled control. Following labeling, cells were immediately lysed with Radio-Immunoprecipitation Assay (RIPA) lysis buffer (Invent, IN-WB001). The volume of the RIPA buffer (100 or 200 μL) was adjusted according to the cell number. Following a 20-min incubation on ice, the mixture was supplemented with DNase (5 or 7 μL, corresponding to the lysis volume) and incubated at 37°C for 20 min, followed by addition of Proteinase K (5 or 7 μL) and a further 10-min incubation at 37°C. Finally, 1 mL RNAiso Plus was added to the lysate immediately for RNA isolation. RNA directly isolated from labeled cells served as untreated control. The isolated total RNA was treated again with DNase and Proteinase K to remove residual genomic DNA and protein. The purified RNA was then precipitated and analyzed by qRT-PCR as described above.

**Cell viability assay using Cell Counting Kit-8 (CCK-8)**

HEK293T cells were seeded at a comparable density and cultured in PDL-coated 96-well plates. Cells were then labeled with HoeDBF according to the labeling protocol for the depletion-based method. The labeling conditions comprised a full factorial combination of HoeDBF concentrations (0, 2.5, 5, and 10 μM) and green light irradiation durations (0, 2, and 3 min). Following the labeling, cells were washed once with DBPS and 100 μL culture medium containing 10% (v/v) enhanced CCK-8 solution (Beyotime, C0046) was added to each well. After incubation at 37°C for 30 min, the absorbance of each well at 450 nm was detected using a microplate reader.

**U1 AMO inhibition**

AMOs were ordered from Gene Tools. The U1 AMO (GGTATCTCCCCTGCCAGGTAAGTAT) [2] was designed to be complementary to the 5'-end recognition sequence of U1 snRNA to inhibit its function. A standard control AMO (CCTCTTACCTCAGTTACAATTTATA) [2] provided by Gene Tools was used as a negative control. HEK293T cells were cultured in standard culture medium at 37 ºC under 5% $CO_2$ and nucleofected upon reaching 70–80% confluence. The transfection was performed using the 4D-Nucleofector (Lonza Bioscience) with program DH-135 [3]. To perform nucleofection for fluorescence confocal imaging and RNA-sequencing, 1 x

$10^6$ cells were transfected with 25 μM U1 or control AMO (2.5 nmol AMO in 100 μL transfection solution) using the SF Cell Line 4D-Nucleofector X Kit L (Lonza Bioscience, V4XC-2024). In contrast, to perform nucleofection for U1 AMO concentration titration, 2 x $10^5$ cells were transfected with U1 AMO at different specified concentrations using SF Cell Line 4D-Nucleofector X Kit S (Lonza Bioscience, V4XC-2032). Following transfection, cells were transferred to a PDL-coated 6-well plate containing warm medium and cultured at 37°C for 4 hours. The cells were then used for subsequent experiments.

**RNase H protection assay followed by qRT-PCR analysis**

To validate U1 blocking efficiency, we performed RNase H protection assay followed by qRT-PCR analysis, which was adapted from a previously described method [4]. The underlying principle of this method is illustrated in Fig. S3A.

**RNase H protection assay using total RNA:** A total of 15 μg RNA was first incubated in a 23.5 μL mixture containing 3 μM U1/scrambled AMO and 1 μL RNaseOUT in nuclease-free $H_2O$ at 37°C for 30 min. Subsequently, 3 μL ASO stock, 0.5 μL RNase H, and 3 μL 10x RNase H buffer were added to the mixture and the resulting 30 μL mixture (containing 3 μM ASO, 2.5 U RNase H, and 1x RNase H buffer) was incubated at 50°C for 20 min. For the negative controls shown in Fig. S3B, the respective components that needed to be omitted were substituted with nuclease-free $H_2O$. Following the incubation, 36 μL nuclease-free $H_2O$, 1.5 μL DNase, and 7.5 μL 10x DNase buffer were added and the resulting mixture was incubated at 37°C for 30 min to digest the excess ASO. The RNA was then isolated with RNAiso Plus and treated with proteinase K to remove residual protein contamination. The purified RNA was then precipitated and analyzed using qRT-PCR.

**RNase H protection assay for U1 AMO concentration titration:** The transfected cells were labeled with HoeDBF according to the labeling protocol for the depletion-based method (labeling conditions: 5 μM HoeDBF incubation, 2-min green light irradiation). The cells were then lysed using 10 μL RIPA lysis buffer supplemented with 1 μL RNaseOUT. The cell lysates were transferred to 1.5 mL conical tubes and incubated on ice for 30 min. Next, 4 μL master mix containing ASO, RNase H and RNase H buffer was added to each tube, resulting in a 15 μL reaction mixture containing 1 μM ASO, 0.25 U RNase H, and 1x RNase H buffer. The reaction mixture was then incubated at 37°C for 30 min. Following the incubation, RNA was isolated using RNAiso Plus for subsequent qRT-PCR analysis.

**Library preparation and high-throughput RNA sequencing**

For HoeDBF-mediated, enrichment-based method, approximately 100 ng of enriched and input RNA was used for library preparation. Ribosomal RNAs (rRNAs) were first removed from the samples using NEBNext rRNA Depletion Kit v2 (Human/Mouse/Rat) (NEB, E7400), following the manufacturer's instructions. The resulting rRNA-depleted RNA was then used for library preparation using NEBNext® Ultra™ II Directional RNA Library Prep Kit (NEB, E7760), following the manufacturer's instructions. Briefly, fragmentation was performed by incubating the samples in a mixture of NEBNext First Strand Synthesis Reaction Buffer and Random Primers at 94°C for 8 min. Following fragmentation, first strand cDNA was synthesized directly in the same mixture by adding NEBNext Strand Specificity Reagent and NEBNext First Strand Synthesis Enzyme Mix, followed by sequential incubation at 25°C for 10 min, 42°C for 50 min, and 70°C for 15 min. The second strand cDNA was then synthesized by adding NEBNext Second Strand Synthesis Reaction Buffer with dUTP Mix and NEBNext Second Strand Synthesis Enzyme Mix, followed by incubation at 16°C for 1 h. The synthesized cDNA products were purified using SPRIselect Beads (Beckman Coulter, A63881). Following end preparation for the cDNAs, adapter ligation was performed. During this step, the second strand of the cDNAs were digested with USER Enzyme. The resulting first strand cDNA library was then purified through a second round of selection using SPRIselect beads. To enrich for cDNA libraries with an insert size larger than 200 bp, two rounds of size selection were performed using SPRIselect Beads (0.3x followed by 0.15x). The selected libraries were then amplified for 13–14 cycles by PCR, and the final products were purified with 0.8x SPRIselect Beads. The size distribution of final cDNA libraries was in the range of 300–500 bp. The cDNA libraries were sequenced on an Illumina Hiseq X 10 platform, yielding approximately 33 million paired-end reads (2x 150 bp) per library.

For PCRD-seq, total RNA was extracted from HEK293T cells (including those subjected to various labeling conditions and those underwent U1 or control nucleofection) with RNAiso Plus, followed by DNase and Proteinase K treatment. The purified RNA samples were sent to GENEWIZ (Suzhou) for stranded RNA library construction and subsequent high-throughput sequencing. Each library yielded approximately 33 million paired-end reads.

For PCRD-seq profiling of RNA from nuclear lamina, mitochondria, endomembrane, lysosomes, and stress granules, the cell lysates within RNAiso Plus of the samples were subjected to RNA isolation, DNase and Proteinase K treatment, stranded RNA library construction and high-throughput sequencing by GENEWIZ (Suzhou). Each library generated 40–60 million paired-end reads.

For PCRD-seq profiling of RNA from HM and NM cells, RNA isolation, purification, as well as library preparation were performed in our own laboratory. Specifically, cells were labeled with HoeDBF according to the labeling protocol for the depletion-based method (labeling conditions: 5 μM HoeDBF incubation, 2-min green light irradiation). The labeled cells were lysed immediately by RNAiso Plus for RNA isolation. The RNA was then treated with DNase and Proteinase K. For library construction, approximately 300 ng of the purified total RNA was used, following a protocol identical to that employed in the enrichment-based method. The size distribution of final cDNA libraries was in the range of 300–500 bp. The cDNA libraries were sequenced on an Illumina NovaSeq X plus platform, yielding approximately 33 million paired-end reads (2x 150 bp) per library.

**Chromatin-associated RNA identification**

Adaptors and low-quality bases were first removed from raw reads using Trimmomatic (v.0.39) [5]. The clean reads were then pseudoaligned against the human reference transcriptome (GRCh38.p14.v47, GENCODE) using kallisto (v0.50.0) [6] for transcript abundance quantification. The transcript-level abundance estimates were summarized to gene-level using tximport [7], followed by differential expression analysis using DESeq2 [8]. caRNAs were operationally defined as transcripts showing significant up-regulation in enriched samples versus input ($\log_2$(fold change) $\geq$ 1, FDR $<$ 0.05) in the enrichment-based method, or significant down-regulation in positively labeled samples versus negative controls ($\log_2$(fold change) $\leq$ -1, FDR $<$ 0.05) in the depletion-based method. The RNA-seq data of chromatin fractionation [9] was downloaded and reanalyzed as described above for comparison. The caRNAs were defined as transcripts exhibiting significant higher abundance ($\log_2$(fold change) $\geq$ 1, FDR $<$ 0.05) in chromatin pellet extract compared to soluble-nuclear extract.

**Unspliced/spliced ratio calculation**

The unspliced/spliced ratio was calculated as described [10]. Briefly, a custom FASTA file was generated by modifying the human reference transcriptome (GRCh38.p14.v47, GENCODE) to include both spliced and unspliced versions for each transcript. This modified transcriptome served as the reference for quantifying transcript abundance with kallisto (v0.50.0) [6], enabling competitive assignment of reads to either spliced or unspliced transcripts. Transcript-level abundances were then summarized to gene level using tximport [7]. Transcripts for which both the spliced and unspliced versions had a TPM $\geq$ 0.1 were retained for subsequent analysis. The unspliced/spliced ratio was defined as (TPM of unspliced transcript) / (TPM of spliced transcript).

For depletion-based method, the calculated unspliced/spliced ratios for enriched and input samples were presented as a density plot (Fig. 3E). For enrichment-based method, the average unspliced/spliced ratio of three enriched sample replicates was divided by the average ratio of the corresponding input samples. The resulting fold-change values were $\log_2$-transformed and visualized as a density plot (Fig. 2D).

**Analysis related to 5’ splice site prediction**

The prediction of 5’ SS was performed as previously described [11, 12] with some modifications. The 9 bp sequences spanning positions - 3 to + 6 relative to exon-intron junctions (i.e., 3 bp in the exon and 6 bp in the intron) and containing a conserved GU dinucleotide at positions 4 and 5 were defined and extracted as 5’ SS [13]. The extracted sequences were then converted to 5’ SS motif using sites2meme. Subsequently, fimo was used to find significant 5’ SS motifs ($p < 0.0001$) in each isoform of the caRNAs. In the 5’ SS strength analysis, to determine the strength of the detected motifs, they were scored by a Maximum Entropy model [14]. Motifs with scores above and below the median of the population were classified as “strong” and “weak”, respectively. A transcript was considered to possess a strong or weak 5’ SS as long as one of its isoforms contained one. In the 5’ SS frequency analysis, the transcripts were divided into quartiles based on the frequency of predicted 5’ SS motifs per kilobase. The 5’ SS frequency of a transcript was calculated as the average frequency of all its isoforms. Only lncRNAs were used for the 5’ SS strength and frequency analysis.
